# Single-cell multimodal profiling of proteins and chromatin accessibility using PHAGE-ATAC

**DOI:** 10.1101/2020.10.01.322420

**Authors:** Evgenij Fiskin, Caleb A Lareau, Gökcen Eraslan, Leif S Ludwig, Aviv Regev

## Abstract

Multi-modal measurements of single cell profiles are a powerful tool for characterizing cell states and regulatory mechanisms. While current methods allow profiling of RNA along with either chromatin or protein levels, connecting chromatin state to protein levels remains a barrier. Here, we developed PHAGE-ATAC, a method that uses engineered camelid single-domain antibody (‘nanobody’)-displaying phages for simultaneous single-cell measurement of surface proteins, chromatin accessibility profiles, and mtDNA-based clonal tracing through a massively parallel droplet-based assay of single-cell transposase-accessible chromatin with sequencing (ATAC-seq). We demonstrate PHAGE-ATAC for multimodal analysis in primary human immune cells and for sample multiplexing. Finally, we construct a synthetic high-complexity phage library for selection of novel antigen-specific nanobodies that bind cells of particular molecular profiles, opening a new avenue for protein detection, cell characterization and screening with single-cell genomics.

---

Massively-parallel single-cell profiling has become an invaluable tool for the characterization of cells by their transcriptome or epigenome, deciphering gene regulation mechanisms, and dissecting cellular ecosystems in complex tissues (Klein et al., 2015; Lareau et al., 2019; Macosko et al., 2015; Satpathy et al., 2019). In particular, recent advances have highlighted the power of multimodal single-cell assays (Ma et al., 2020), such as cellular indexing of transcriptomes and epitopes by sequencing (CITE-seq), that profile both transcriptome and proteins by DNA-barcoded antibodies (Mimitou et al., 2019; Peterson et al., 2017; Stoeckius et al., 2017).

Although the vast combinatorial space of oligonucleotide barcodes theoretically allows parallel quantification of an unrestricted number of epitopes, in practice, however, we are limited by the availability of antigen-specific antibodies. Moreover, each antibody must be separately conjugated with a unique oligonucleotide (oligo)-barcode, which currently does not allow a scalable and pooled construction of barcoded antibody libraries. Finally, technologies for the combined high-throughput measurement of the epigenome and proteome have not been described.

To overcome these limitations, we developed PHAGE-ATAC (**Figures 1A-1C, Supp. Fig. 1**), a multimodal single-cell approach for phage-based multiplex protein measurements and chromatin accessibility profiling using droplet-based scATAC-seq (10x Genomics scATAC (Satpathy et al., 2019)). PHAGE-ATAC enables sensitive quantification of epigenome and proteins, captures mtDNA that can be used as a native clonal tracer (Lareau et al., 2020; Ludwig et al., 2019), introduces phages as renewable and cost-effective reagents for high-throughput single-cell epitope profiling, and leverages phage libraries for the selection of antigen-specific antibodies (Hoogenboom, 2005; Smith, 1985), altogether providing a novel platform that greatly expands the scope of the single-cell profiling toolbox.

**Figure 1.**
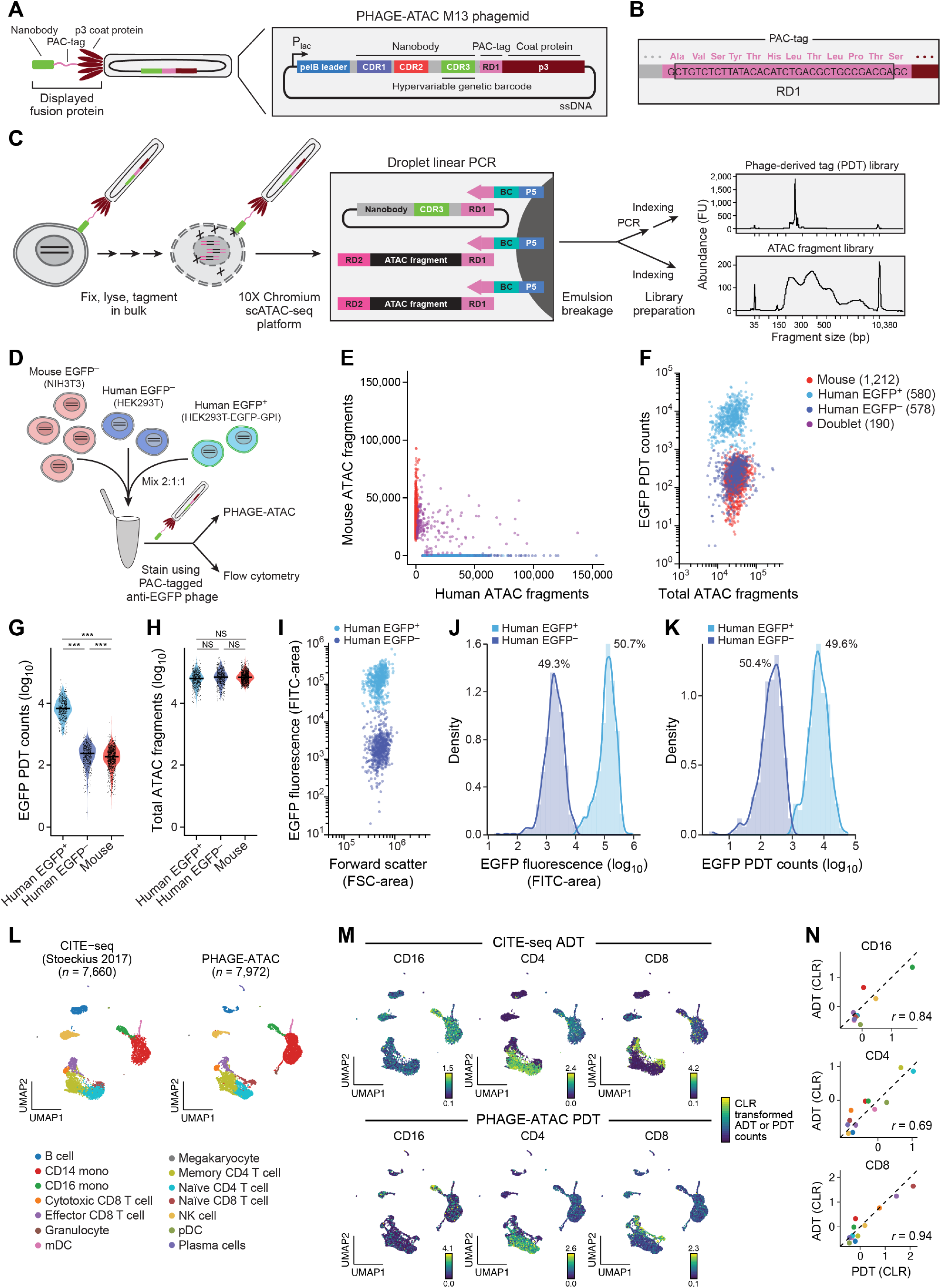
PHAGE-ATAC for massively-parallel simultaneous measurement of protein epitopes and chromatin accessibility. **(A)-(C)** PHAGE-ATAC overview. **(A)** Schematic of engineered nanobody-displaying M13 phages used for PHAGE-ATAC. Nanobodies are displayed via fusion to p3, the PAC-tag is placed in the linker between nanobody and p3. M13 phagemids contain a pelB leader for periplasmic secretion and incorporation of fusions during phage assembly. **(B)** PAC-tag RD1 sequence (pink) allows capture by 10x ATAC gel bead oligos (**Supp. Fig. 2A**), without interrupting translation (right). **(C)** PHAGE-ATAC workflow. After phage nanobody staining, fixation, lysis and tagmentation in bulk (leftmost), single cells and 10x ATAC gel beads are encapsulated into droplets using 10x microfluidics, followed by linear amplification with simultaneous droplet barcoding of chromatin fragments and phagemids via hybridization of 10x barcoding primers to RD1 sequences (second from left). Separate PDT and ATAC sequencing libraries are prepared (second from right and **Supp. Fig. 3**). Right: Representative BioAnalyzer traces. BC, bead barcode. **(D)-(K)** Single-cell ATAC and EGFP specificity in a species-mixing experiment. **(D)** Experimental scheme. **(E)** Number of human (x axis) and mouse (y axis) ATAC fragments associated with each bead barcode (dots), colored by assignment as human EGFP+ (light blue), human EGFP- (dark blue), mouse (red), doublet (purple, >10% human and mouse fragments). **(F)** EGFP PDT counts (y axis, log_10_ scale) and number of ATAC fragments (x axis, log_10_ scale) for each bead barcode (dots) colored as in E (color legend). **(G),(H)** Distributions of EGFP PDTs (G, y axis, log_10_ scale) and ATAC fragments (H, y axis, log_10_ scale) in each of the three populations (x axis) (Mann-Whitney one-tailed, ***p < 10^−4^, NS=not significant). Line: median. **(I)-(K)** PDT quantification is consistent with flow cytometry. EGFP fluorescence (I, y axis) and distribution (J, x axis) and distribution of EGFP PDT (K, x axis) in EGFP+ (light blue) and EGFP- (dark blue) human cells. **(L)-(N)** PHAGE-ATAC and CITE-seq compare well in human PBMCs. **(L),(M)** Two-dimensional joint embedding of scRNA-seq profiles from PBMCs from published CITE-seq (Stoeckius et al., 2017) and of scATAC-seq profiles from PBMCs generated by PHAGE-ATAC, colored by annotated cell types (L) or by the level of protein marker ADTs (M, top) or PDTs (M, bottom). **(N)** Agreement between protein level estimates from CITE-seq and PHAGE-ATAC. ADT (y axis, centered log ratio (CLR)) and PDT (x axis, CLR) for each marker gene across cell types (dots, colored as in **L**), Pearson’s *r* is shown.

Protein quantification in PHAGE-ATAC is based on epitope recognition by nanobody (Ingram et al., 2018) (Nb)-displaying phages (**Figure 1A**, **Supp. Fig. 1**), in contrast to recognition by oligonucleotide-conjugated antibodies in CITE-seq and related methods (Peterson et al., 2017; Stoeckius et al., 2017), or fluorescently labeled antibodies in other techniques (Katzenelenbogen et al., 2020; Paul et al., 2015). The hypervariable complementarity-determining region 3 (CDR3) within each Nb-encoding phagemid acts as a unique genetic barcode (Pollock et al., 2018) that is identified by sequencing in PHAGE-ATAC and serves as a proxy for antigen detection and quantification (**Figure 1A**, **Supp. Fig. 1A**). To allow phage-based epitope quantification alongside accessible chromatin using droplet-based scATAC-seq, we engineered an M13 phagemid for the in-frame expression of (**1**) an epitope-binding Nb, (**2**) a PHAGE-ATAC tag (PAC-tag) containing the Illumina Read 1 sequence (RD1) and (**3**) the phage coat protein p3 for surface display (**Figures 1A and 1B**). This enables phage Nb (pNb)-based recognition of cell surface antigens, simultaneous droplet-based indexing of phagemids and ATAC fragments, as well as separate generation of phage-derived tag (PDT) and ATAC sequencing libraries (**Figure 1C, Supp. Fig. 2 and 3, Methods**).

We first confirmed that the PHAGE-ATAC modified phagemid workflow allows successful and specific pNb antigen recognition and pNb-based cell staining during scATAC cell lysis. As a first proof-of-concept, we used HEK293T cells expressing surface-exposed glycosylphosphatidylinositol (GPI)-anchored EGFP (EGFP-GPI) that are specifically recognized by an anti-EGFP pNb (Rothbauer et al., 2006) (**Supp. Fig. 4A-4E**). Importantly, introducing the PAC-tag did not impair Nb display and antigen recognition (**Supp. Fig. 4F and 4G**). Moreover, fixation retained pNb-based cell staining after the scATAC lysis step, with a standard scATAC-seq buffer (**Supp. Fig. 5, Methods**).

To benchmark PHAGE-ATAC for single cell profiling, we performed a ‘species-mixing’ experiment, in which we pooled mouse (NIH3T3), human EGFP^-^ (HEK293T) and human EGFP^+^ (HEK293T-EGFP-GPI) cells at a 2:1:1 ratio, followed by anti-EGFP pNb staining, library generation and analysis using a custom computational workflow (**Figure 1D and Supp. Fig. 6; Methods**). After filtering, we recovered 1,212 mouse and 1,158 human cell barcodes (**Figure 1E**), with good library complexity, enrichment of fragments in peaks, and enrichment in transcription start sites (**Supp. Fig. 7A-7C**), all comparable to gold-standard published reference data without additional protein detection (Lareau et al., 2020; Satpathy et al., 2019). Analysis of EGFP PDT counts confirmed the presence of EGFP^+^ and EGFP^-^ cells (**Figures 1F and 1G**) that together with mouse cell barcodes were all recovered at expected input ratios (observed 2.09:1:1, expected 2:1:1), with no substantial differences in scATAC-seq data quality metrics (**Figure 1H, Supp. Fig. 7**). EGFP PDT levels by PHAGE-ATAC (**Figures 1F and 1G**) and EGFP fluorescence intensities by standard flow cytometry (**Figure 1I**) were highly concordant (**Figures 1J and 1K**). Taken together, these results established the use of PDTs for accurate and sensitive epitope quantification in single cells concomitantly with scATAC-seq.

Next, we showed that PHAGE-ATAC can discern cellular states of primary peripheral blood mononuclear cells (PBMCs) comparably to CITE-seq. For PHAGE-ATAC, we targeted well-characterized markers via a panel of three pNbs targeting CD4, CD8 and CD16 using previously reported high-affinity Nb sequences (Roobrouck et al., 2016; Tavernier et al., 2017), as well as anti-EGFP as a negative control (**Methods**). Flow cytometry of pNb-stained PBMCs and side-by-side comparison between pNb and conventional antibody-stained cells confirmed the antigen-specificity of the produced phages (**Supp. Fig. 8**). In addition, we further optimized the PHAGE-ATAC lysis buffer to better preserve phage staining (Lareau et al., 2020) (**Supp. Fig. 9; Methods**). Integrative canonical correlation analysis (Butler et al., 2018), clustering and dimensionality reduction of PHAGE-ATAC data of 7,972 high-quality PBMCs and published CITE-seq data of 7,660 PBMCs (Stoeckius et al., 2017) (**Figure 1L, Methods**) identified the same set of expected cell states and markers (**Figure 1L and Supp. Fig. 10A**). The distribution of PDTs and CITE-seq antibody-derived tags (ADTs) across all cell types were highly correlated for each surface marker (**Figures 1M and 1N**, Pearson’s *r* = 0.69-0.94). To further validate PDT partitioning independently of CITE-seq, we determined differential gene activity scores from the PHAGE-ATAC data alone by comparing scATAC profiles of T cells based on CD4 and CD8 PDT abundances (**Supp. Fig. 10B and 10C**). This identified both *CD4* and *CD8* loci as top hits and recovered many known *bona fide* markers of CD4^+^ and CD8^+^ T cells (*e.g*. CD4: *CTLA4, CD40LG, ANKRD55*; CD8: *PRF1, EOMES, RUNX3*, **Supp. Fig. 10C**). Finally, EGFP PDTs were only detected at background levels, confirming the high specificity of pNbs (**Supp. Fig. 10D and 10E**). These results illustrate the capacity of PHAGE-ATAC to reliably and specifically detect endogenous cell surface proteins in single cells along with their epigenomic profiles.

To scale PHAGE-ATAC, we next introduced a cost-effective alternative for sample multiplexing in scATAC-seq using pNbs for Cell Hashing. A number of current methods allow ‘overloading’ antibody-tagged cells into droplets to increase single-cell processing throughput and mitigate batch effects (Gehring et al., 2020; Lareau et al., 2019; McGinnis et al., 2019; Stoeckius et al., 2018). To demonstrate hashtags for PHAGE-ATAC, we generated four anti-CD8 hashtag pNbs (henceforth referred to as hashtags) by introducing different silent mutations into the anti-CD8 CDR3 (**Figure 2A, Methods**), allowing sequencing-based identification of the four hashtags. As expected, the hashtags displayed comparable CD8 recognition within PBMCs (**Supp. Fig. 11A**). To demonstrate phage-based hashing, we stained CD8 T cells from each of four healthy donors with a unique hashtag, pooled them and processed the pool by PHAGE-ATAC, overloading 20,000 cells (**Figure 2A**) (*vs*. ~6000 cells without overloading). These yielded high-quality data for 8,366 cell barcodes, to which we assigned donor and singlet/doublet status from hashtag counts (**Methods**), identifying the sample of origin for 6,438 singlets and 703 doublets (observed doublet rate 8.4% compared to 10% expected) (**Figure 2B**). As expected, barcodes assigned to an individual hashtag had higher count distributions for the respective hashtag (**Figure 2C**). Singlet and doublet assignments were concordant with a two-dimensional embedding of hashtag count data (**Figure 2D**), with the expected higher numbers of chromatin fragments and hashtag counts in doublets (p < 2.2×10^−16^; Mann-Whitney test, **Figures 2E and 2F**). The hashtag-based assignments were also highly concordant with assignments based on computationally derived donor genotypes from accessible chromatin profiles (Heaton et al., 2020) (**Methods**), with a singlet classification accuracy of 99.3% and an overall classification accuracy of 92.9% (**Figure 2G**). Interestingly, chromatin accessibility analyses revealed a small set of putative B cells (**Supp. Fig. 11B and 11C**), consistent with the presence of a minor contaminating population after CD8 T cell enrichment. While B cells were classified as hashtag-negative, genotype and hashtag-based classification were highly consistent across CD8 T cell states (**Figure 2H and Supp. Fig. 11D-11F**), confirming hashtag antigen specificity.

**Figure 2.**
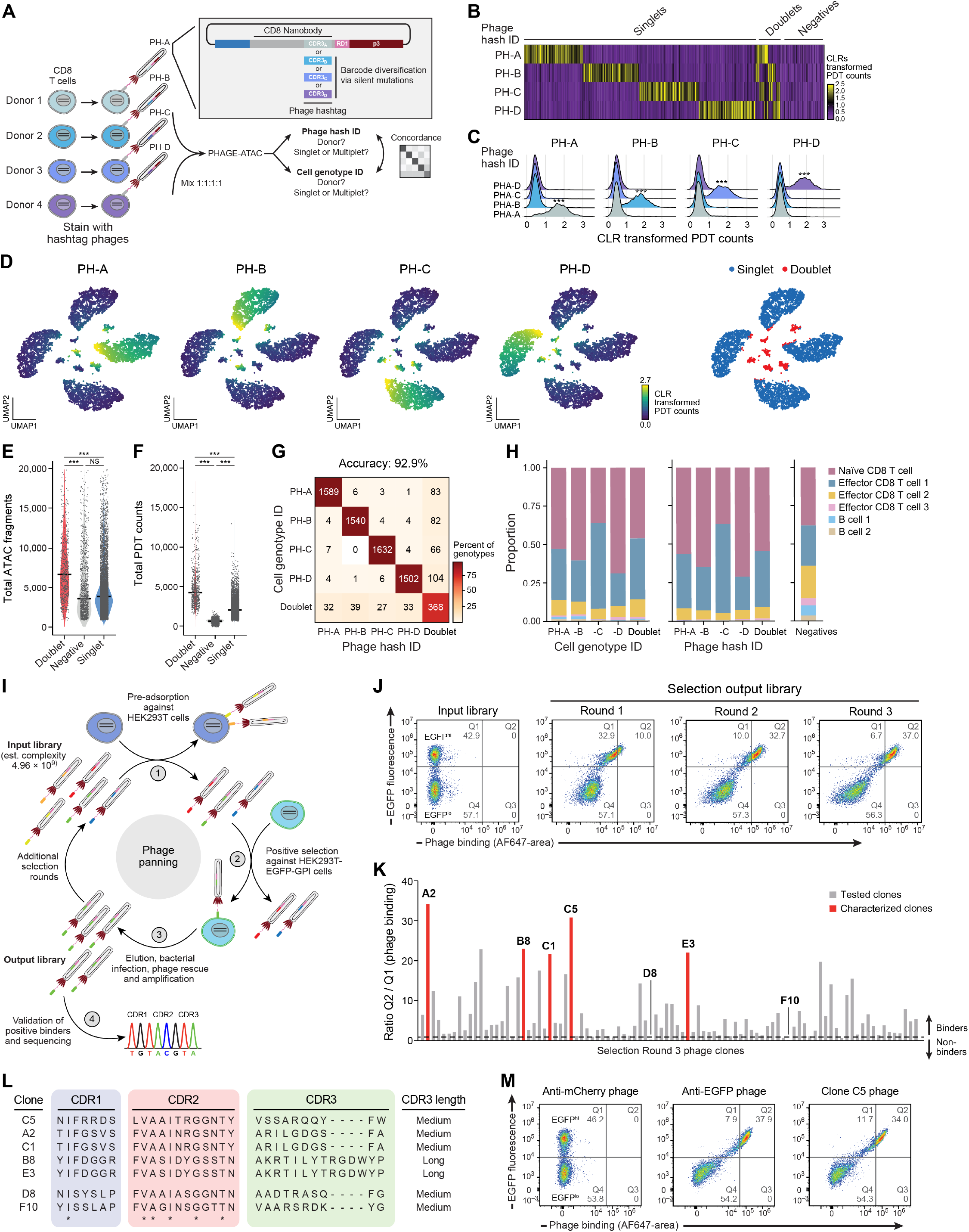
PHAGE-ATAC compatible phage nanobodies enable sample multiplexing and can be selected using phage display. **(A)** Generation of phage hashtags by silent mutations. Shown is a schematic for four anti-CD8 phage hashtags and a subsequent hashing experiment using CD8 T cells from four human donors. **(B)-(H)** Effective demultiplexing of phage hashtags. (**B**) PDT counts (color bar, CLR) for each hashtag (rows) across cells (columns) sorted by their HTODemux classification (Phage hash ID). (**C**) PDT count distributions for each hashtag (colored histograms) across the four Phage hash IDs (Wilcoxon two-tailed, ***p < 10^−4^). (**D**) Two-dimensional embedding of cell barcodes by PDT count data, colored by PDT count for the marked hashtag (4 left panels) or by singlet/doublet classification (right). (**E**), **(F)** Distribution of the number of ATAC fragments per barcode (E, y axis) or PDT counts (F, y axis) in cell barcodes in each category (x axis) (Mann-Whitney two-tailed, ***p < 10^−4^, NS=not significant). Line: median. (**G)** Number and percent (color) of barcodes shared between each genotype-based (Genotype ID, rows) and Phage hashtag ID-based (columns) assignments. Top: overall accuracy. (**H**) Proportion of cells of each type (y axis) within each assigned barcode category (x axis) based on either genotype (left) or and hashtags (right), and in the negative fraction (far right). **(I)-(M)**, Selection of PHAGE-ATAC nanobodies by phage display. (**I**) Schematic of phage display selection using PANL (**Methods**). PANL is panned against EGFP-expressing cells (HEK293T-EGFP-GPI) with preceding counter-selection against antigen-devoid parental cells (HEK293T). Bound phages are eluted, used to infect bacterial hosts and output libraries are generated. After multiple selection rounds, antigen-recognizing phage nanobody clones are picked, phagemids are isolated and nanobody inserts are sequenced. (**J**) Flow cytometry analysis of selection progress. Flow cytometry plots of EGFP fluorescence (y axis) and phage binding (x axis, AlexaFluor647 area) to EGFP-GPI-expressing HEK293T cells (EGFP^hi^ and EGFP^lo^) in, from left, the input library and after each of three consecutive selection cycles (see also **Supp. Fig. 4C and Methods**). **(K)** Flow cytometry screen of 94 phage nanobody clones derived from selection round 3. Ratio of Q2 to Q1 signal (as defined in **Figure 1J**) when staining EGFP-GPI-expressing HEK293T (EGFP^hi^ and EGFP^lo^) cells with individual phage nanobodies after the 3^rd^ round of selection. Dashed line: threshold of Q2/Q1=1 used for calling positive clones. **(L)** CDR sequences and CDR3 length of selected clones obtained by Sanger sequencing. * non-randomized constant positions in PANL library (see also **Supp. Fig. 12A**). **(M)** Flow cytometry plots of EGFP fluorescence (y axis) and phage binding (x axis, AlexaFluor647 area) to EGFP-GPI-expressing HEK293T cells (EGFP^hi^ and EGFP^lo^) using an immunization-based (Rothbauer et al., 2006) anti-EGFP Nb-displaying phage (middle), clone C5 from our screen (right) and an anti-mCherry phage negative control (left).

PHAGE-ATAC also enables the concomitant capture of mitochondrial genotypes via mitochondrial DNA-derived Tn5 fragments (Lareau et al., 2020), providing a third data modality that relates protein and accessible chromatin profiles to cell clones. Mitochondrial genotyping using mgatk (Lareau et al., 2020) was broadly concordant with the hashtag assignments, but showed that two donors (PH-B and PH-C) had indistinguishable mitochondrial haplotypes, whereas each of the other two donors had several distinguishing mitochondrial variants (**Supp. Fig. 11G**). Collectively, these results established the use of hashtag pNbs for sample multiplexing in scATAC-seq, and its ability to capture mtDNA for clonal analysis.

The production of novel high-quality antigen-specific antibodies is laborious, expensive and limited by animal immunization, generating a bottleneck for antibody-based protein profiling. In contrast, recombinant antibody technology based on phage display has allowed fast and cost-effective selection of high-affinity binders (Miersch and Sidhu, 2012). To enable rapid generation of novel antigen-specific pNbs for PHAGE-ATAC, we developed PHAGE-ATAC Nanobody Library (PANL), a synthetic high-complexity (4.96×10^9^) pNb library (**Supp. Fig. 12**). To demonstrate identification of novel pNbs using PANL, we performed a selection against EGFP-GPI-expressing HEK293T cells, while counter-selecting using parental HEK293T (**Figure 2I**). Over three selection rounds, we monitored the enrichment of pNbs by staining EGFP-GPI^+^ cells, revealing a steady increase of antigen-recognizing pNbs with each additional round (**Figure 2J**). Screening of 94 clones after the final (third) selection demonstrated that at least 95% of clones recognized EGFP-GPI^+^ cells with strong binding (Q2/Q1 >1) (**Figure 2K and Supp. Fig. 13**). As clones varied in their ability to bind EGFP-GPI^+^ cells, we picked 7 clones (5 strong and 2 weak binders) and sequenced their phagemid inserts. Sanger sequencing uncovered the presence of multiple identical clones (A2 and C1, B8 and E3, **Figure 2L**), illustrating selection-driven convergence. Finally, side-by-side comparison of a selected clone (C5) and a reported high-affinity anti-EGFP Nb derived from immunized animals (Rothbauer et al., 2006) indicated similar binding to EGFP-GPI^+^ cells (**Figure 2M**). These results demonstrate the utility of PANL for the rapid selection of pNbs to detect and quantify antigens of interest on cells. They further illustrate PANL’s potential for the generation of a new toolbox of barcoded affinity reagents for single cell genomics.

In conclusion, PHAGE-ATAC uses the power of recombinant phage display technology as the basis for single cell profiling of cell surface proteins, chromatin accessibility and mtDNA. This allows users to leverage the renewable nature, low cost and scalability of pooled phage library preparation as well as the compact size and stability of nanobodies (Ingram et al., 2018). We envisage PHAGE-ATAC as an adaptive tool that may be further combined with unique molecular identifiers for phagemid counting and other engineerable scaffolds used in phage display applications (e.g. scFv, Fab) (Gebauer and Skerra, 2009). In the future, we believe this will significantly enhance our ability for the cost-effective (**Supp. Fig. 14**), multimodal single-cell characterization of the proteome, epigenome and likely additional readouts at an unprecedented depth and specificity.

## Supplementary Figure legends

**Supplementary Figure 1:**
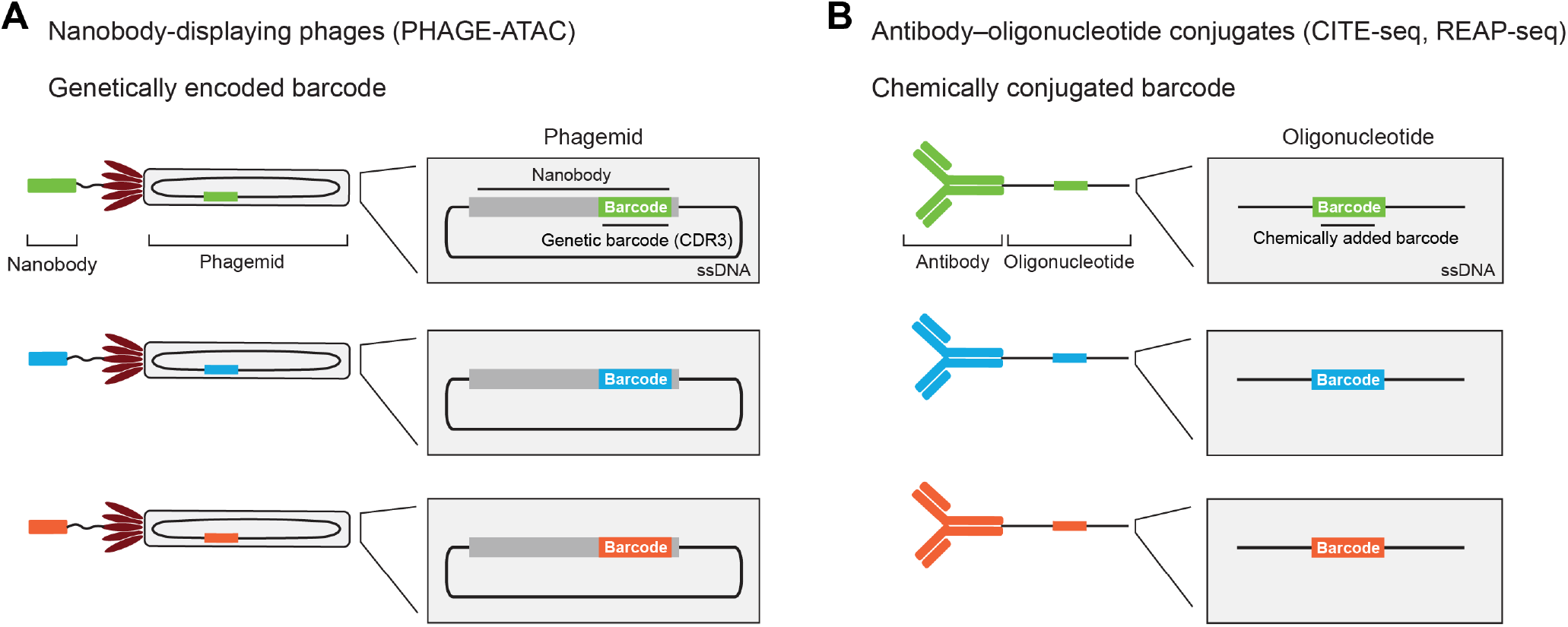
Barcoding strategies for epitope quantification by PHAGE-ATAC vs. CITE-seq. **(A)** Nanobody-displaying phages for PHAGE-ATAC. The phagemid contained within a particular phage particle encodes the protein displayed on that same phage, and PHAGE-ATAC leverages the hypervariable nanobody CDR3 sequences as unique genetic barcode identifiers for each phage. **(B)** Oligonucleotide-conjugated antibodies for CITE-seq. Each antibody is separately conjugated with a unique DNA-barcode.

**Supplementary Figure 2.**
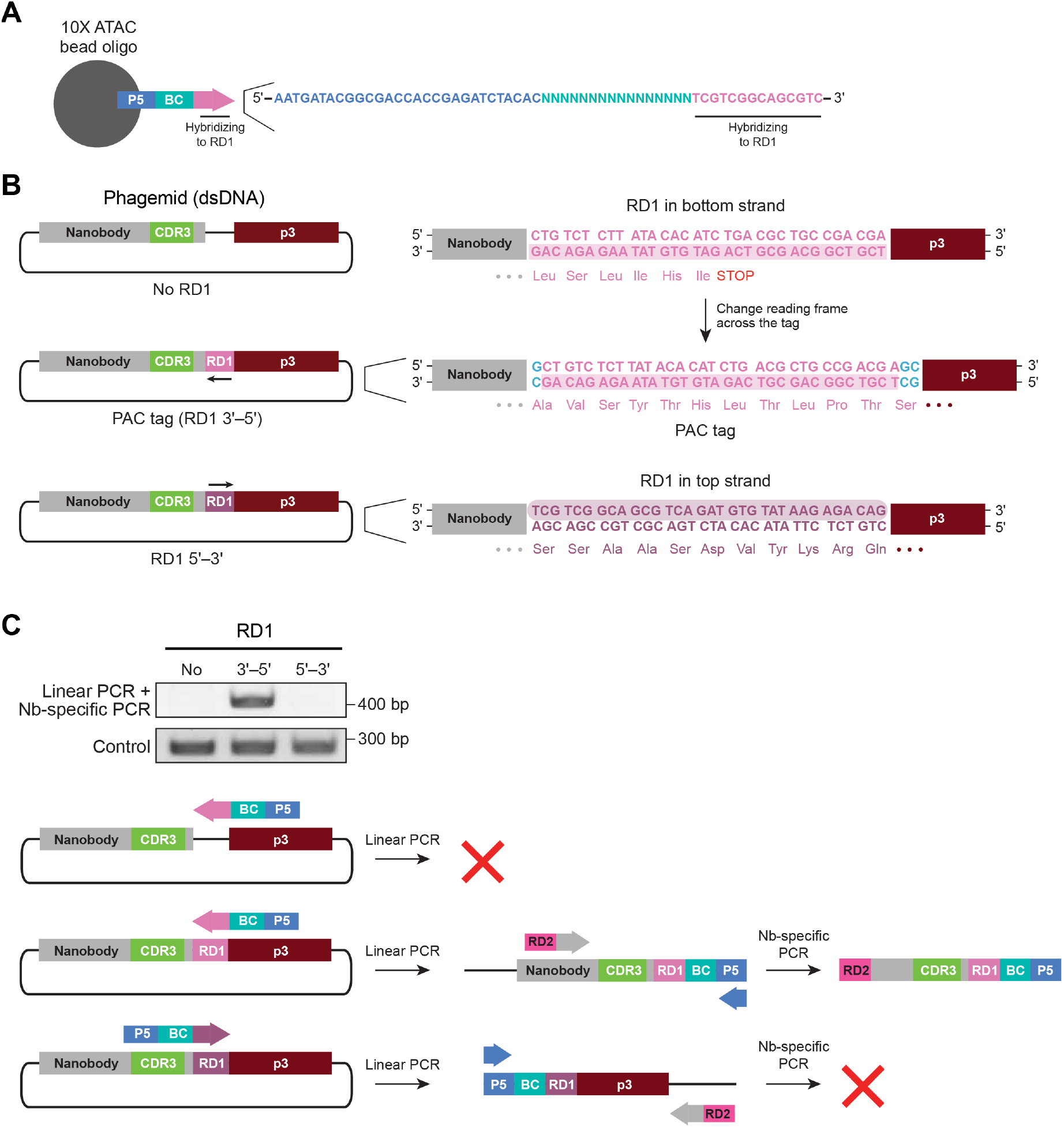
Phage barcode amplification using 10x Genomics scATAC-seq primers enabled by a modified Illumina Read 1 (RD1) sequence. **(A)** Schematic of gel bead oligos showing Illumina P5 sequence (P5), random bead barcode (BC) and the first 14bp of RD1 used for hybridization with RD1-containing chromatin fragments and engineered PHAGE-ATAC phagemids. **(B)** Nanobody-encoding phagemid constructs for RD1-mediated CDR3 barcode capture by 10x Genomics primers. The top strand is the coding strand. Orientation (arrows and shaded boxes), nucleotide sequence and translation product of RD1-containing constructs are shown. To avoid generating a stop codon by introduction of RD1 into the nanobody-p3 reading frame additional codons are introduced to maintain the reading frame across RD1, thus establishing the PAC tag. **(C)** Agarose gel after two-step PCR consisting of linear amplification using the 10x ATAC primer followed by exponential PCR using P5 and Illumina Read 2 (RD2)-containing nanobody-specific primers. PDTs were only obtained for PAC-tagged phagemids with RD1 located on the non-coding strand (3’-5’ orientation relative to nanobody). Abbreviations as in **A**. Control PCR was performed using two primers hybridizing within the nanobody sequence **Methods**).

**Supplementary Figure 3.**
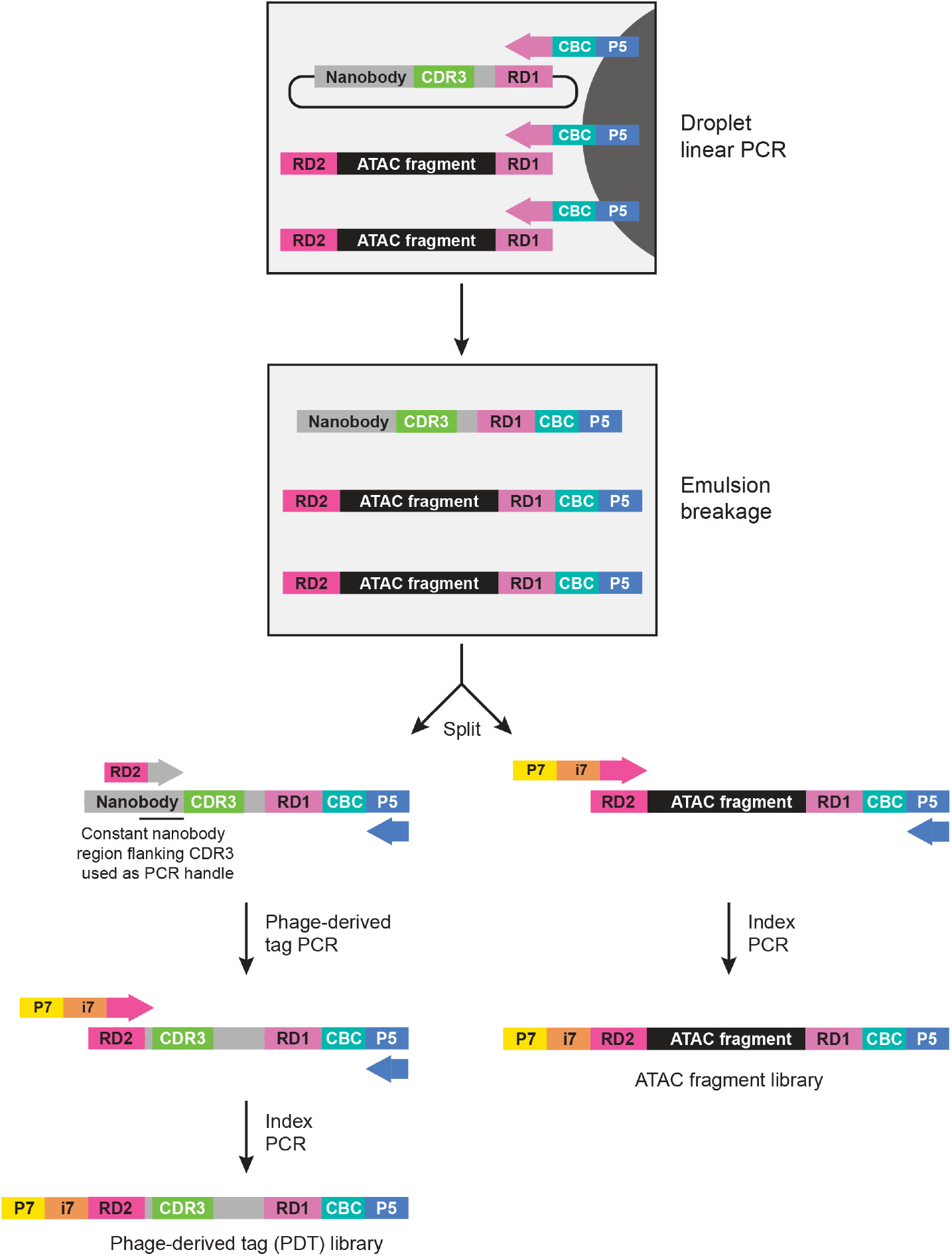
Workflow for separate preparation of scATAC and PDT libraries after droplet-based indexing. Schematic of post barcoding steps for the generation of ATAC and PDT sequencing libraries (**Methods**). After breaking emulsions, barcoded linear amplification products are purified and samples are split. ATAC fragment libraries are immediately processed for sample index PCR. PDT libraries are first amplified in a PDT-specific PCR using a CDR3 flanking constant nanobody sequence as PCR handle. PDT amplification allows RD2 adapter introduction required for final sample indexing. P5 and P7, Illumina P5 and P7 sequences. CBC, random 10x bead cell barcode. i7, sample index.

**Supplementary Figure 4.**
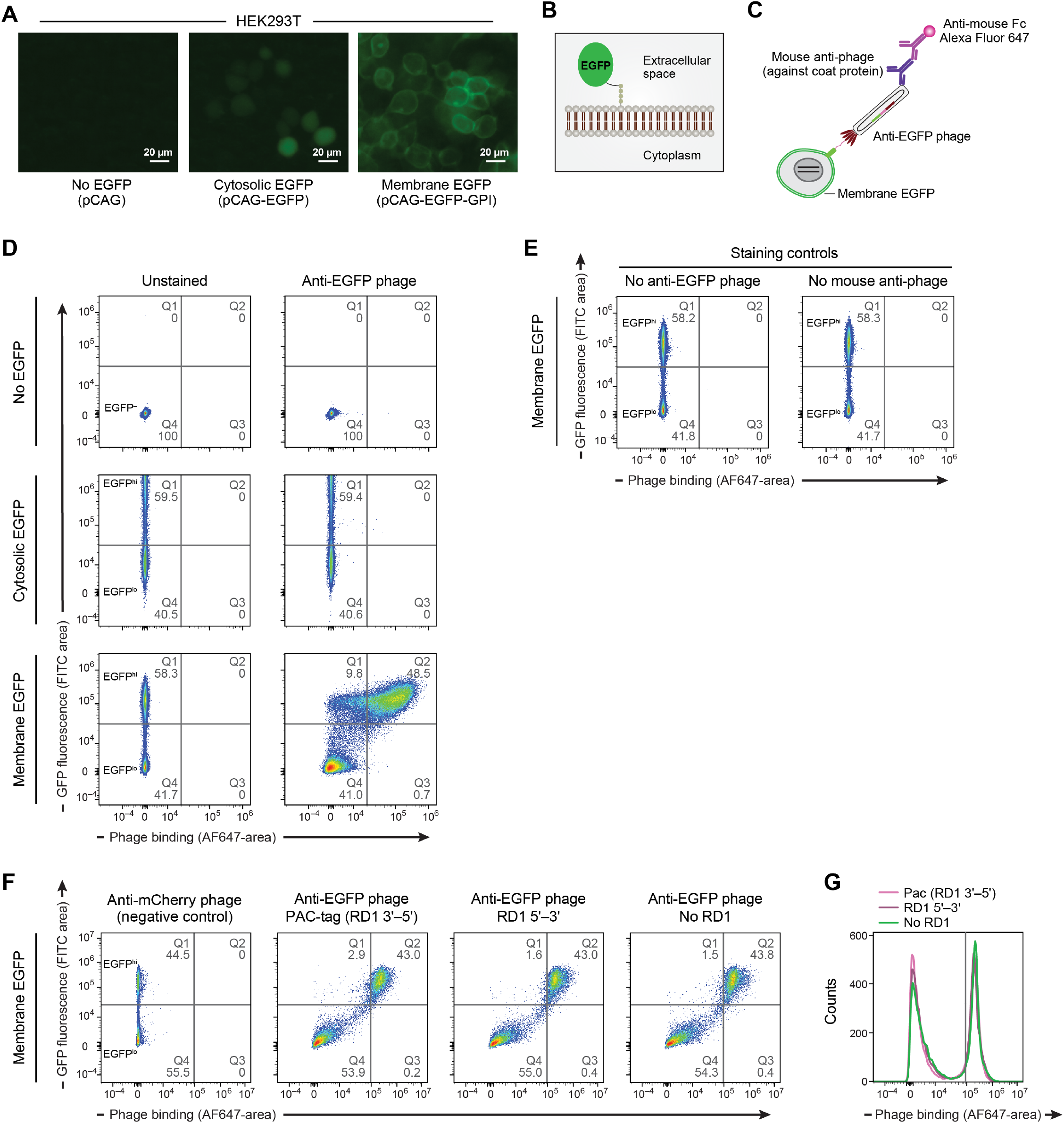
Detection of membrane-localized EGFP via anti-EGFP nanobody-displaying phages. **(A),(B)** Membrane expressed EGFP. **(A)** Microscopy images of HEK293T cells expressing indicated constructs, showing differential localization of untagged cytosolic EGFP (pCAG-EGFP, middle) and GPI-anchored membrane-localized EGFP (pCAG-EGFP-GPI, right, **Methods**). **(B)** Schematic of surface-exposed GPI-anchored EGFP. **(C)** Schematic for detection of phage recognition via flow cytometry. Phage-stained cells are incubated with mouse anti-M13 coat protein antibodies followed by detection by Alexa Fluor 647-conjugated anti-mouse secondary antibodies. Phage binding is thus reflected by Alexa Fluor 647 signal. **(D)** Flow cytometry analysis of anti-EGFP phage nanobody binding to EGFP-expressing HEK293T cells. EGFP fluorescence (y axis) and phage binding (x axis, Alexa Fluor 647) in each of the HEK293T cell populations as in **A**, either unstained (left) or stained with an anti-EGFP phage (right). EGFP-expressing cells were always characterized by the presence of both EGFP^hi^ and EGFP^lo^ populations. **(E)** Specificity of detection. As in **D** but using the indicated staining controls for specific staining of membrane-EGFP-expressing cells. **(F),(G)** PAC-tag does not impact nanobody display and antigen interaction. EGFP fluorescence (F, y axis) and phage binding (F, x axis, Alexa Fluor 647) and distribution of level of phage binding (G) for phage-stained EGFP-GPI expressing cells using indicated phage nanobodies (for RD1 sequences see **Supp. Fig. 2B**).

**Supplementary Figure 5.**
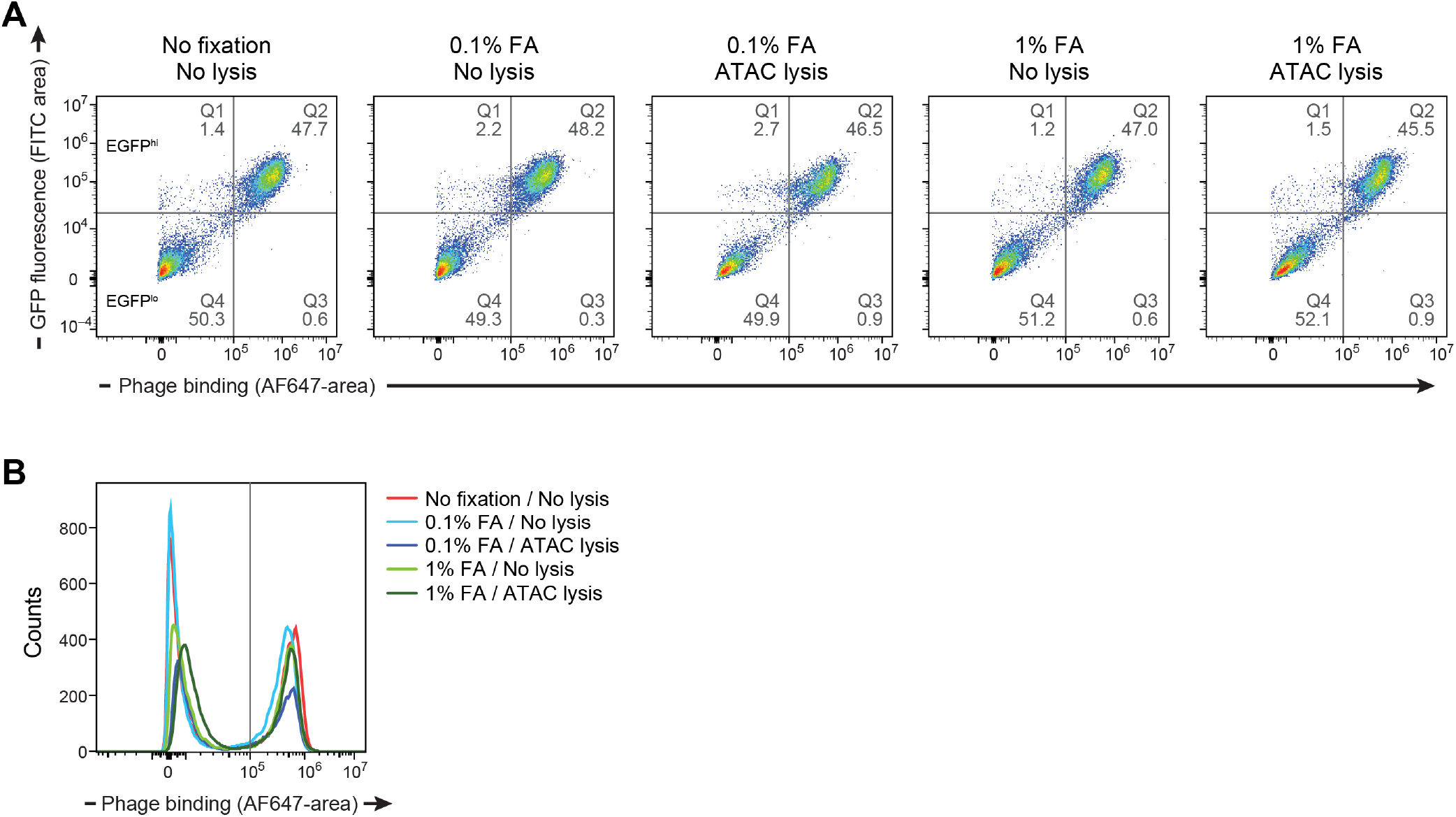
Optimization of fixation and lysis conditions for PHAGE-ATAC species-mixing experiment. EGFP fluorescence (A, y axis) and phage binding (A, x axis, Alexa Fluor 647) and distribution of level of phage binding (B) for EGFP-GPI expressing cells stained with PAC-tagged anti-EGFP-Nb displaying phages after fixation and permeabilization using indicated conditions.

**Supplementary Figure 6.**
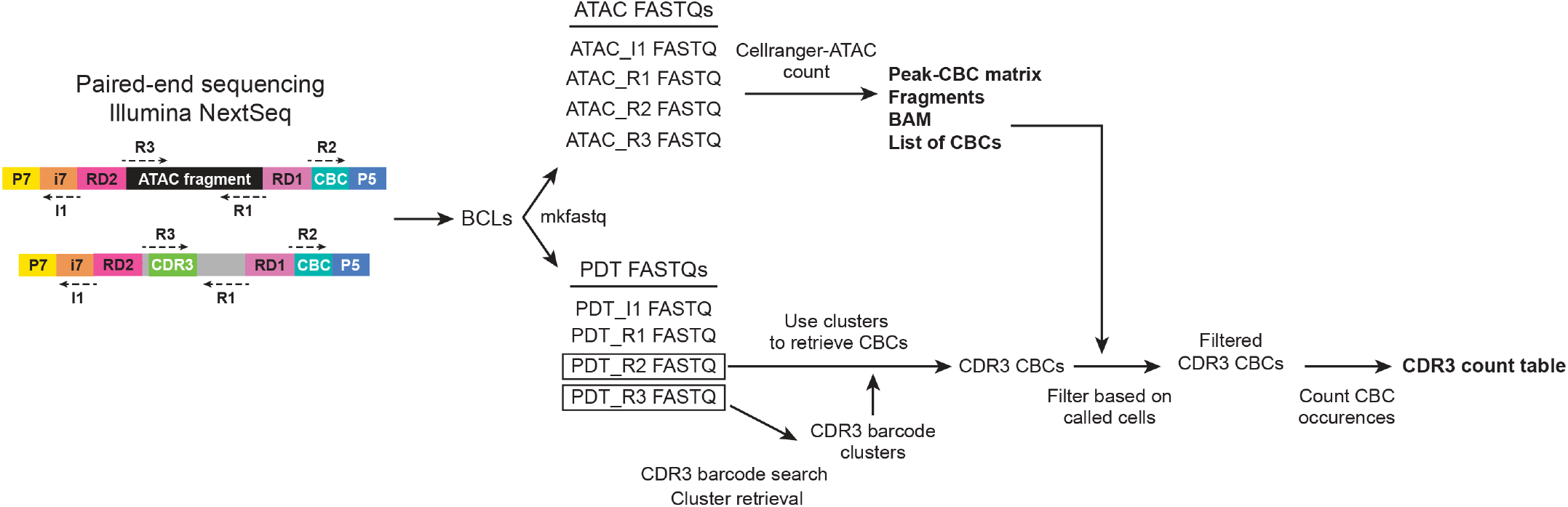
Computational workflow for PHAGE-ATAC data analysis. Paired-end sequencing output is demultiplexed using sample index information (left) to recover ATAC and PDT fastqs. ATAC fastqs are processed using CellRanger-ATAC count for fragment alignment, assignment of cell barcodes and generation of peak-cell barcode matrices. CDR3 barcode sequences are used to search PDT_R3 fastqs and identify CDR3-containing sequencing clusters. Matching of cluster identifiers is used to derive corresponding cell barcodes from PDT_R2 fastqs. Recovered PDT cell barcode lists are filtered using cell barcodes called by CellRanger. Cell barcode occurrences are counted to generate PDT-cell barcode count matrices (**Methods**).

**Supplementary Data Figure 7.**
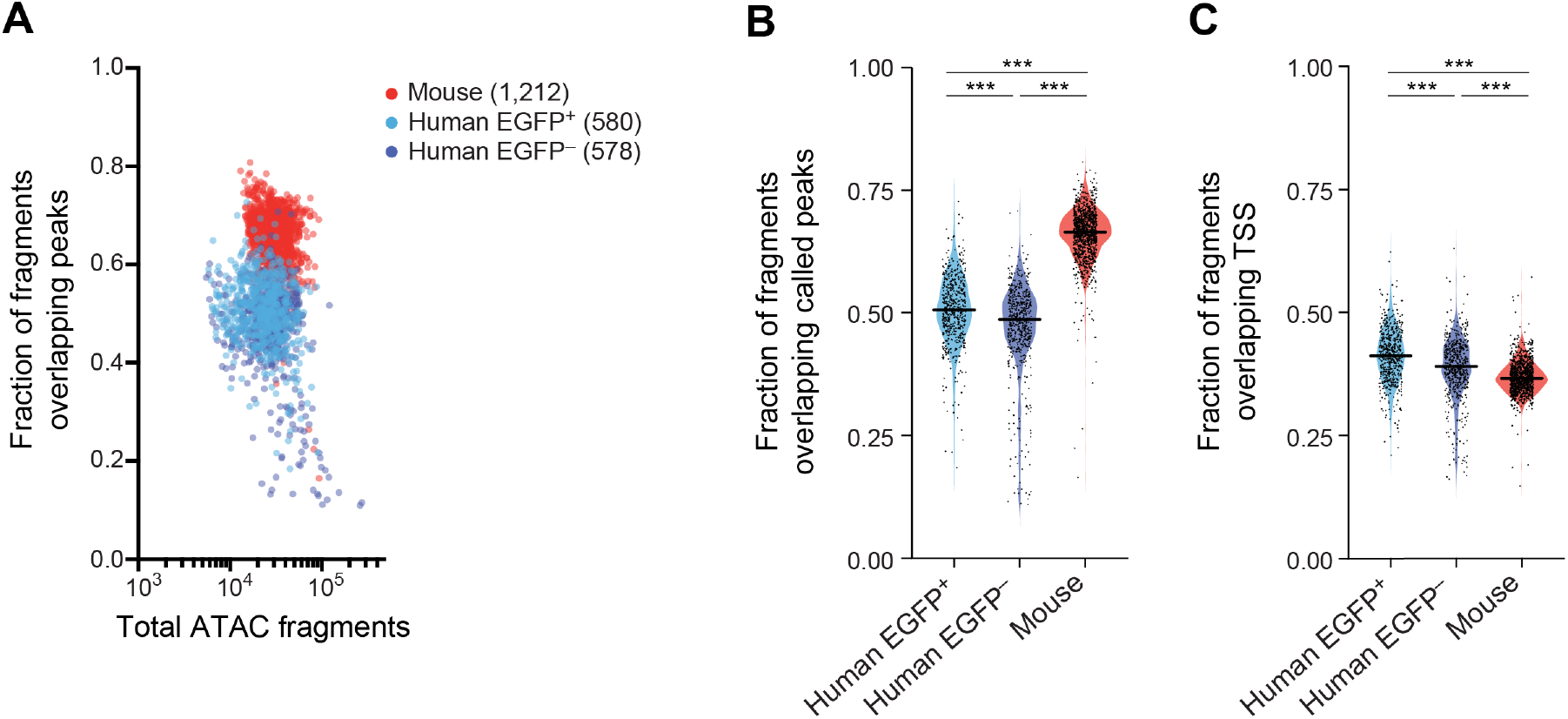
PHAGE-ATAC quality metrics for human-mouse species-mixing experiment. **(A)** Fraction (y axis) and number (x axis, log_10_ scale) of unique chromatin fragments overlapping peaks for each barcode (dot) colored by populations (color legend). **(B),(C)** Distribution of fraction of unique ATAC fragments overlapping peaks (B, y axis) or TSS (C, y axis) in each of the three cell populations (x axis) (Mann-Whitney two-tailed, ***p < 10^−4^, NS=not significant). Line: median.

**Supplementary Figure 8.**
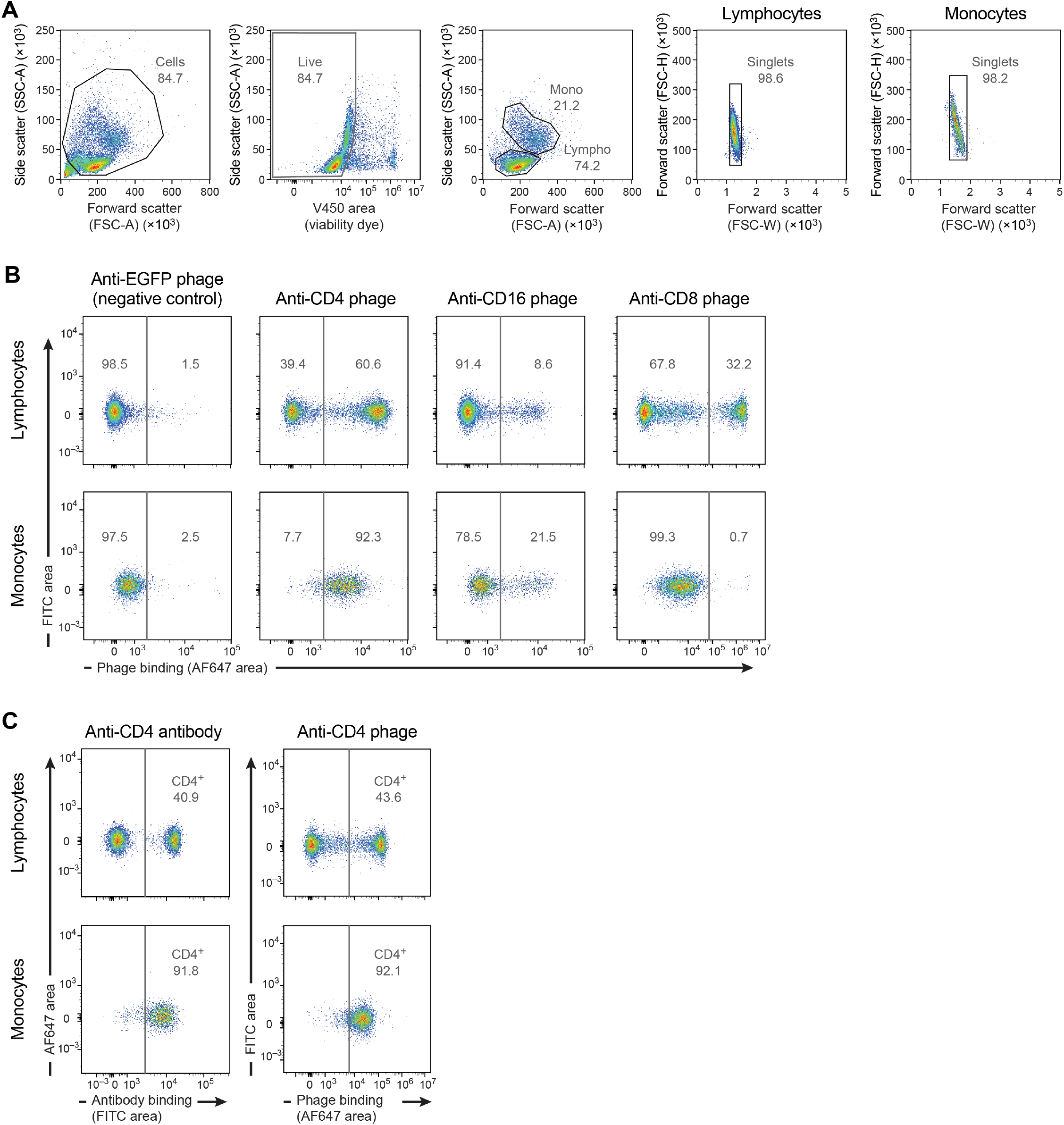
Validation of PAC-tagged anti-CD4, anti-CD8 and anti-CD16 nanobody-displaying phages. **(A)** Flow cytometry gating strategy for analyzed phage-stained PBMCs. **(B)** Flow cytometry-based binding assessment of indicated surface marker-recognizing phage nanobodies to gated lymphocyte and monocyte populations, anti-EGFP pNb was used as negative control. **(C)** Comparison of PBMCs stained with a well-characterized anti-CD4 antibody or generated anti-CD4 phage nanobody. Phage binding is reflected by Alexa Fluor 647 fluorescent signal intensity.

**Supplementary Figure 9.**
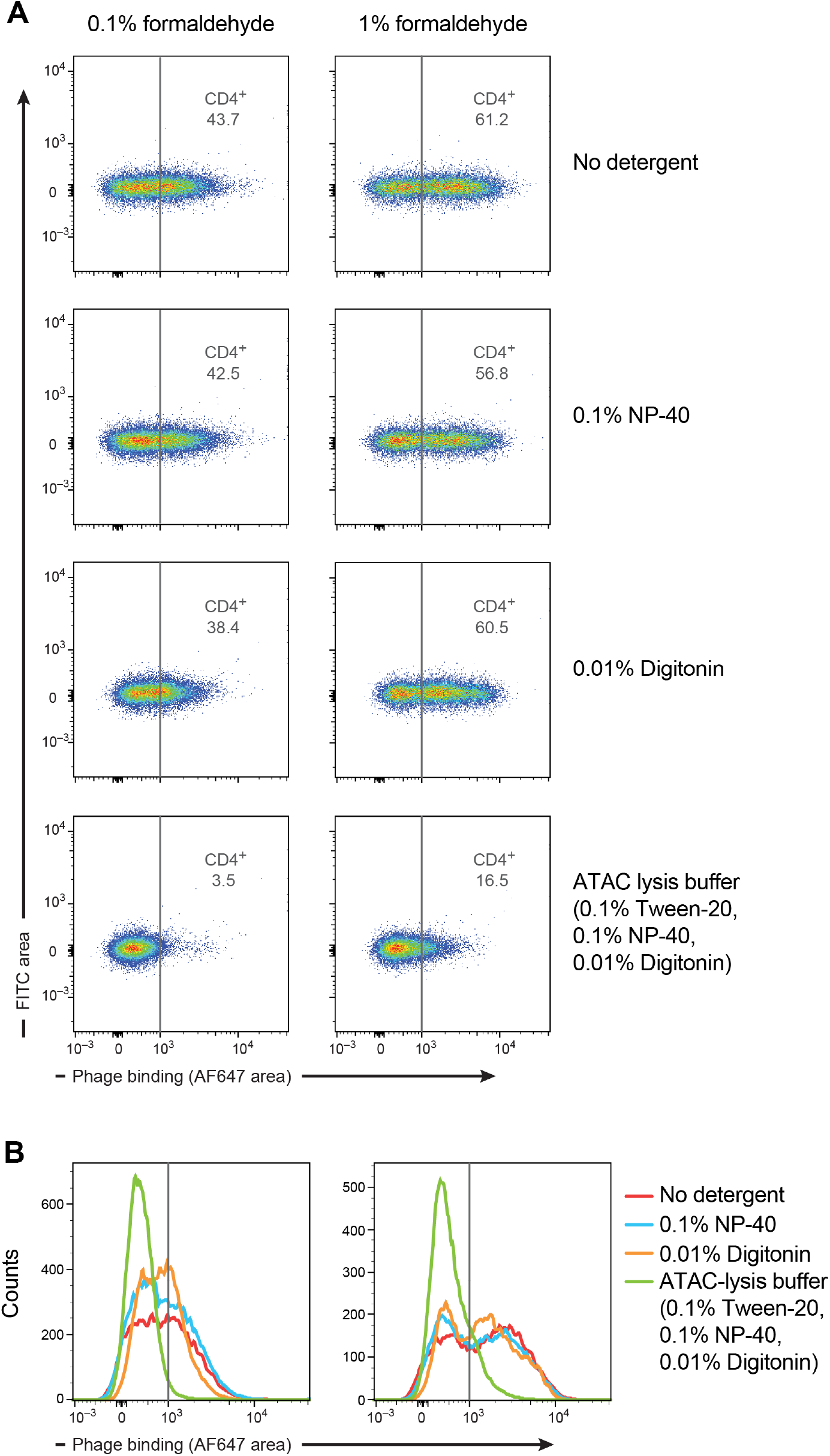
Optimization of fixation and lysis conditions for PHAGE-ATAC using PBMCs. **(A)** Binding of generated anti-CD4 phage nanobodies to PBMCs under indicated conditions. Two different formaldehyde concentrations as well as various depicted lysis buffers were used. Phage binding is reflected by Alexa Fluor 647 fluorescent signal intensity. **(B)** Histogram of data in **(A)**.

**Supplementary Figure 10.**
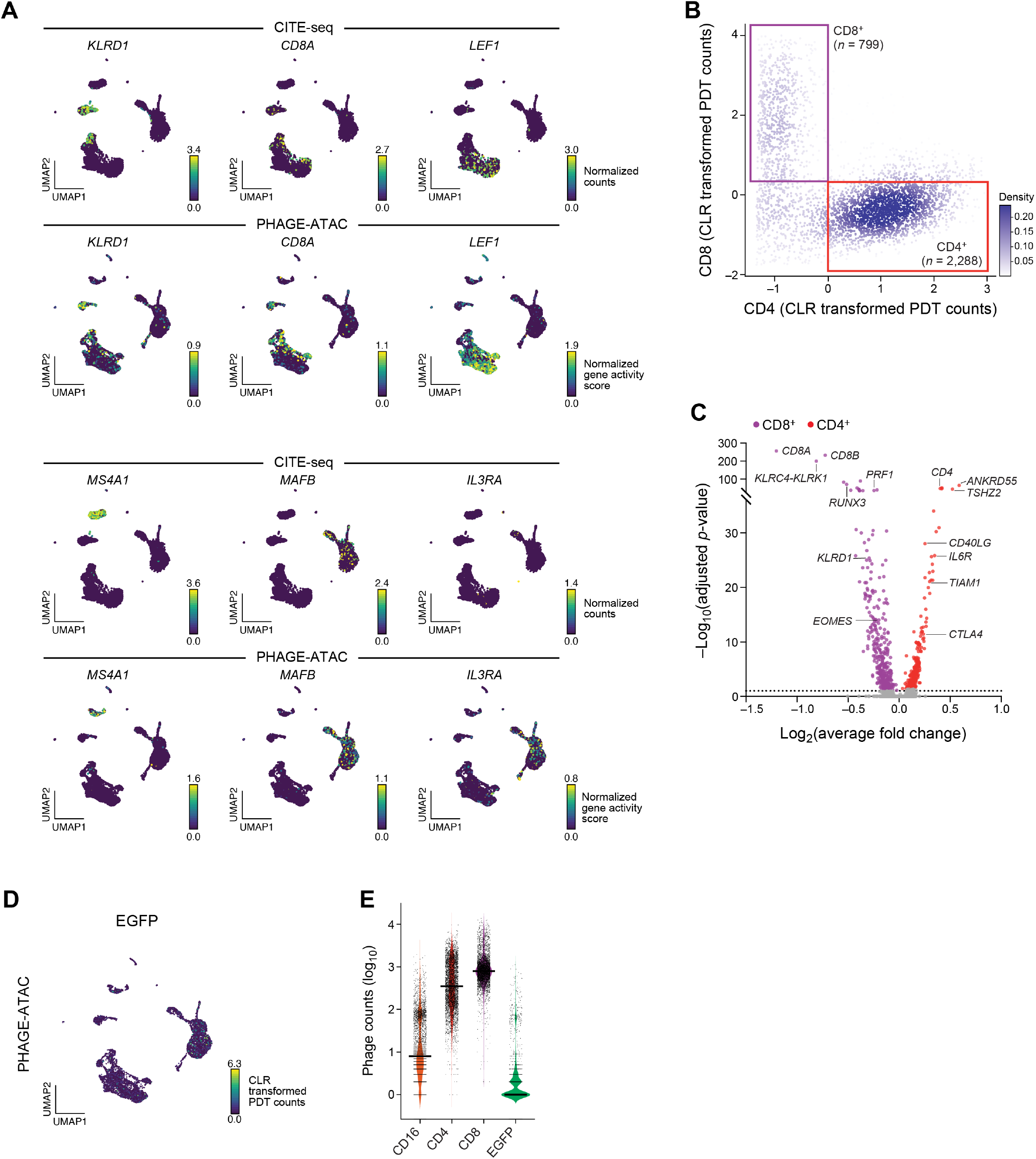
Multimodal single-cell analysis of human PBMCs using PHAGE-ATAC. **(A)** Two-dimensional joint embedding of scRNA-seq profiles from PBMCs from published CITE-seq (Stoeckius et al., 2017) and of scATAC-seq profiles from PBMCs generated by PHAGE-ATAC, colored by the measured RNA level from CITE-Seq (top panels) or by gene activity scores from PHAGE-ATAC (bottom panels) (**Methods**). **(B)**, **(C)** PHAGE-ATAC gating by phage staining highlights cell type specific loci. **(B)** PDT count-based classification of CD4^+^ and CD8^+^ T cells. PDT counts (CLR transformed) of CD8 (y axis) and CD4 (x axis) in each cell (dots). Red boxes: gates for CD4+ and CD8+ cells. **(C)** Average fold change (x axis, log_2_) and associated significance (y axis, −log_10_(P-value) for each gene activity comparing between PDT-classified CD4 and CD8 T cells shown in B. Known *bona fide* markers of either CD4 or CD8 T cells are marked. **(D)** Negative control. Embedding of PHAGE-ATAC data as in A, colored by anti-EGFP pNb PDT. **(E)** Distribution of phage counts (y axis, log_10_) for each cell barcode for each assayed nanobody (x axis).

**Supplementary Figure 11.**
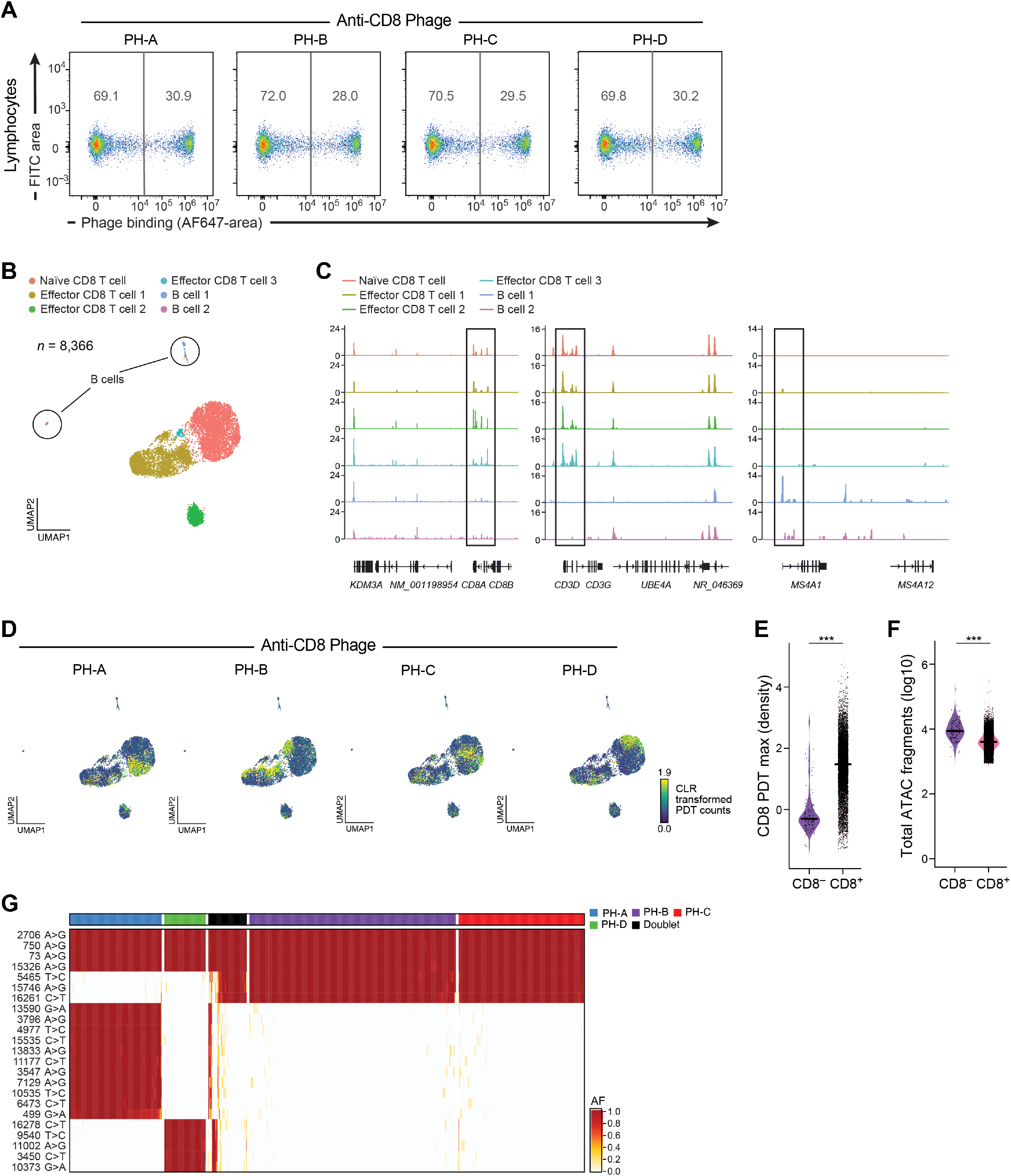
Sample multiplexing using hashtag phages. **(A)** Validation of phage hashtag binding. Flow cytometry of anti-CD8 hashtag phages bound (Alexa Fluor 647 fluorescent signal, x axis) to lymphocytes gated via flow cytometry of phage-stained PBMCs (as shown in **Supp. Fig. 8A**). **(B)** Cell type identification. Two-dimensional embedding of hashed CD8 T cells analyzed by PHAGE-ATAC, colored by cell type annotation. **(C)** Pseudobulk chromatin accessibility track plots for *CD8, CD3* and *MS4A1* (*CD20*) loci across identified cell types. **(D)** Embedding as in B with cells colored by CD8 hashtag PDTs. **(E),(F)** Distribution of maximal CD8 PDT density (E, y axis) or unique chromatin fragments (F, y axis) for each cell barcode in CD8^-^ (B cell 1 and B cell 2) and CD8^+^ (non-B cell) cells (x axis) (Mann-Whitney two-tailed, ***p < 10^−4^). **(G)** Concordance between hashtag-based classification of barcodes and identified mtDNA SNPs. Heteroplasmy (allele frequency percentage; color bar) of different mtDNA variants (rows) in each cell (column), labeled by hashtag assignment (vertical top color bar).

**Supplementary Figure 12.**
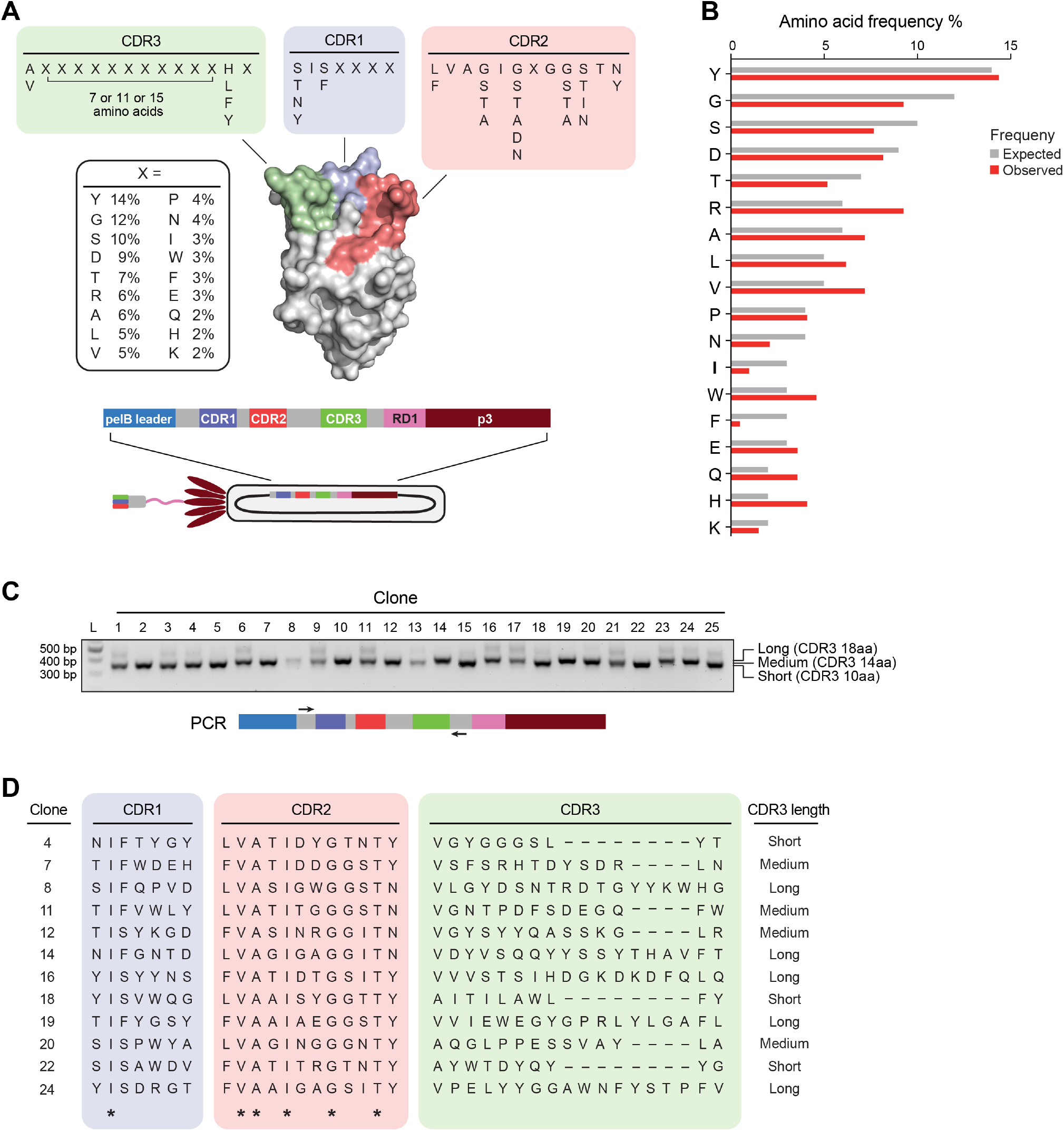
Establishment of PANL, a fully synthetic high-complexity PAC-tagged phage nanobody library. **(A)** Schematic of PANL library design and library phagemid. CDR3 sequence diversification and nanobody framework (grey) in PANL are based on a previously reported nanobody randomization strategy (McMahon et al., 2018). White box: expected frequency of amino acids at each hypervariable position (denoted by X), adjusted by using a custom randomized primer mix for library generation (**Methods**). CDR3 loops contained either 7, 11 or 15 hypervariable positions, resulting in total CDR3 lengths of 10 (short), 14 (medium) or 18 (long) amino acids. Partially randomized positions are depicted as columns, constant positions contain a single amino acid. A deposited structure of anti-EGFP Nb (PDB: 3ogo (Kubala et al., 2010)) with colored CDR3 loops is shown. PANL phagemid is analogous to the one shown in **Figure 1A**. **(B)** Expected (grey) and observed (red) frequencies (x axis) of amino acids at hypervariable positions (y axis) (**Methods**). **(C)** Amplification products of phagemid insert-spanning PCR reactions using depicted primers for 25 randomly picked PANL clones. Product sizes due to presence of long, medium or short CDR3 are shown. **(D)** CDR3 sequences of selected clones from C obtained by Sanger sequencing, CDR3 length is indicated, * non-randomized constant positions in the PANL library.

**Supplementary Figure 13.**
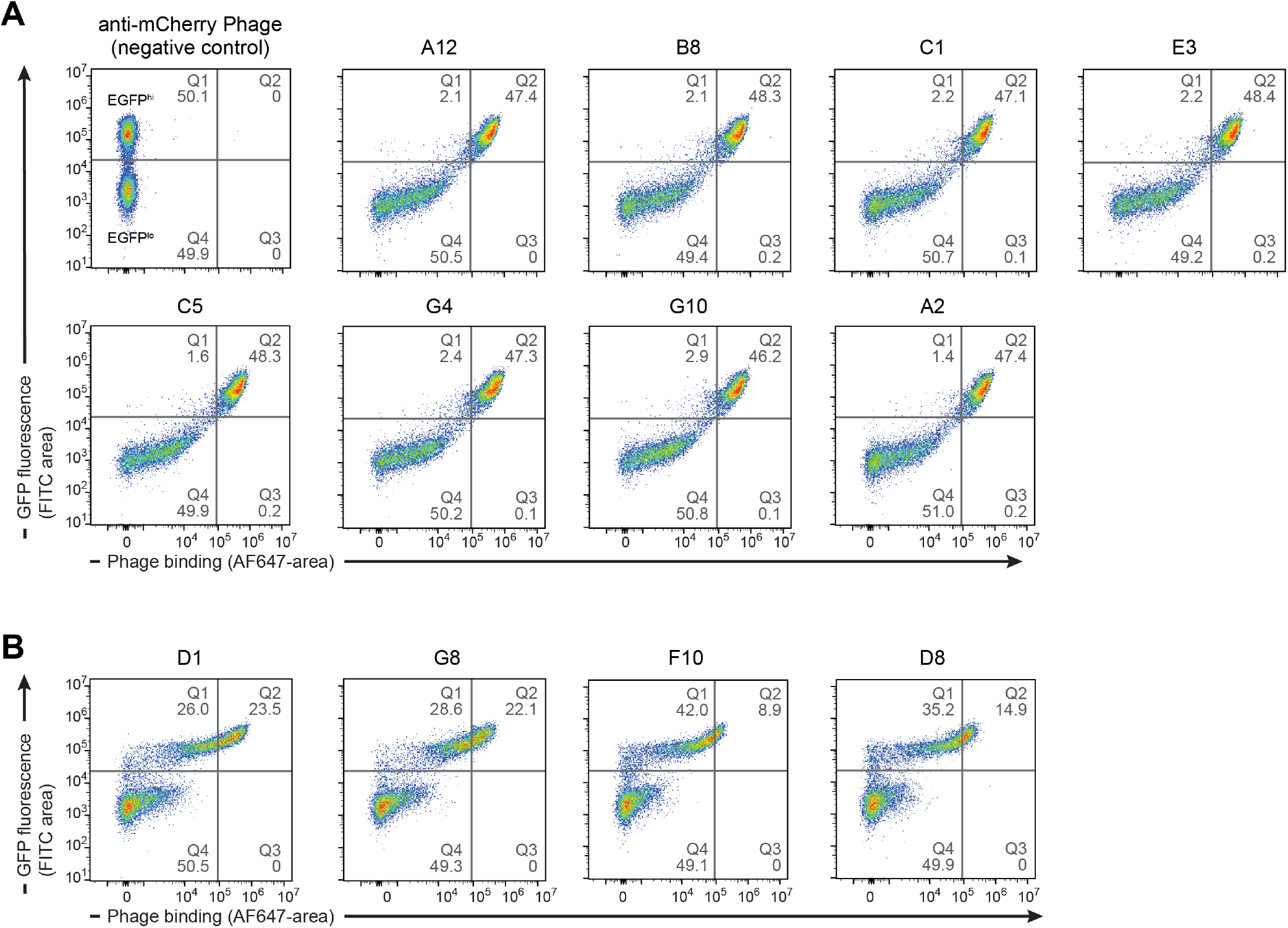
Flow cytometry-based screen of nanobody-displaying phage clones from selection round 3. Flow cytometry analysis of round 3 phage nanobody clones for binding to EGFP-GPI expressing cells (EGFP^hi^ and EGFP^lo^ populations can be observed) with either strong (A) or weaker (B) binders. Phage nanobodies against mCherry were used as negative control. Phage binding is reflected by Alexa Fluor 647 signal.

**Supplementary Figure 14.**
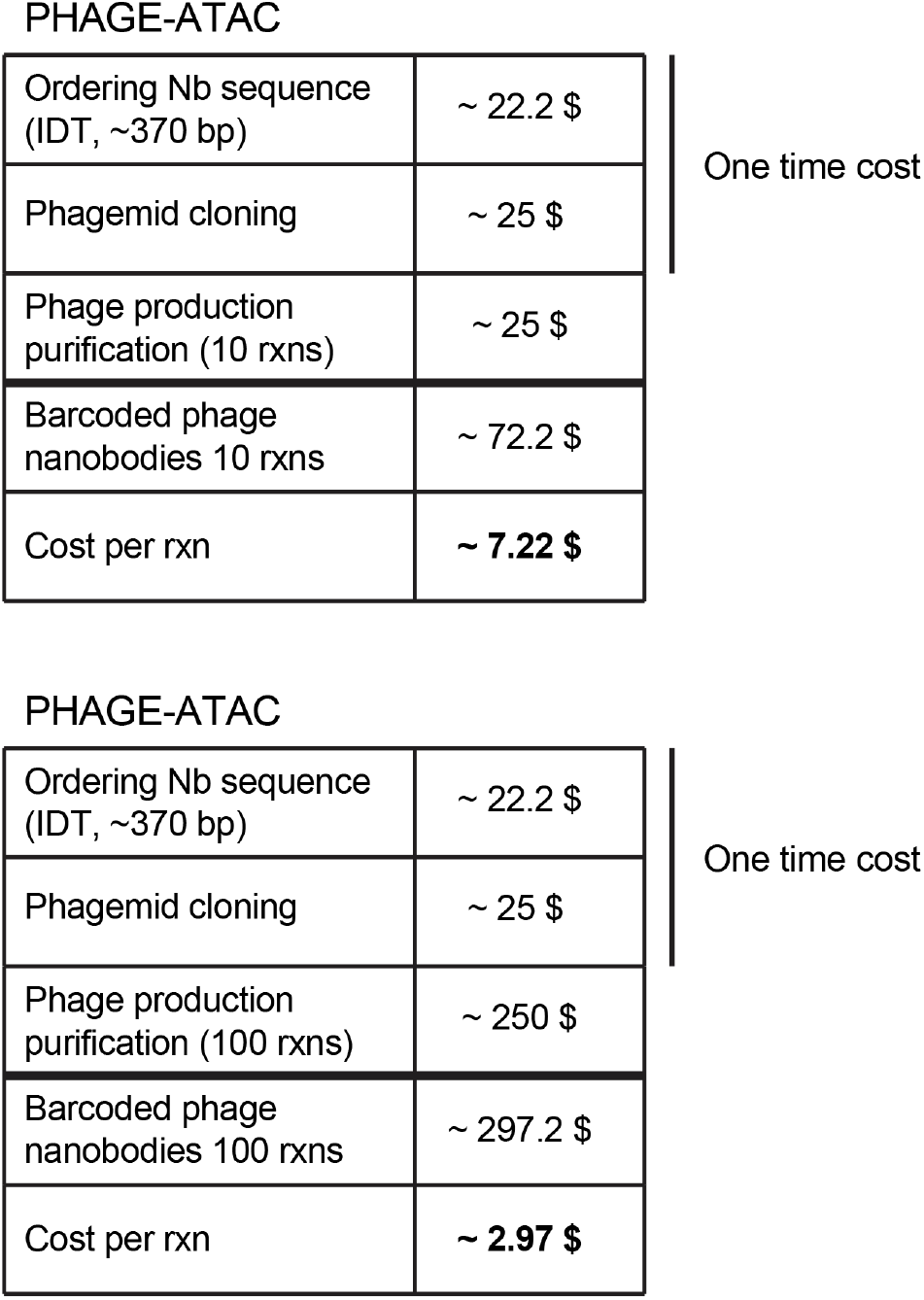
Estimates of cost per reaction for phage nanobodies. Comparison of cost estimates per reaction step and overall for a phage nanobody produced recombinantly.

## Methods

### Oligonucleotides

Oligonucleotide sequences are listed in **Supplementary Table 1**. Oligonucleotides were ordered from Integrated DNA Technologies (IDT) unless indicated otherwise.

### Cloning of phagemids for display of PAC-tagged nanobody-p3 fusions for PHAGE-ATAC

Based on the 10x scATAC bead oligo design (**Supp. Fig. 2A**), we hypothesized that introduction of an RD1 flanking the Nb CDR3 barcode would enable barcode capture alongside accessible chromatin fragments during droplet-based indexing. To avoid premature termination of nanobody-p3 fusion translation due to the introduction of RD1, we modified the RD1-spanning reading frame, which resulted in the expression of a 12-amino acid PHAGE-ATAC tag (PAC-tag). To generate a phagemid for C-terminal fusion of both PAC-tag and p3, 20ng pDXinit (Addgene ID: 110101) were subjected to site-directed mutagenesis with primers EF77 and EF78 using PfuUltraII (Agilent) in 50μl reactions. PCR conditions were 95°C 3min; 19 cycles 95°C 30sec, 60°C 1min, 68° 12min; final extension 72°C 14min. Template DNA was digested for 1.5h at 37°C by addition of 1.5μl DpnI (Fastdigest, Thermo Scientific). PCR reactions were then purified using GeneJet Gel Extraction Kit (Thermo Scientific) and eluted in 45μl water. 20μl eluate were transformed into chemically-competent *E. coli* (NEB Stable Competent) and plated on LB-Ampicillin, yielding pDXinit-PAC. For cloning of nanobody-PAC-p3 fusion-encoding phagemids, nanobody sequences (**Supplementary Table 3**) were ordered as gBlocks from IDT. 25ng nanobody gBlocks were first amplified by PCR to introduce SapI restriction sites. Primers EF87 and EF88 were used for CD4 Nb, primers EF87 and EF89 for CD16 Nb and primers EF104 and EF105 for CD8 Nb. 50μl PCR reactions using Q5 (NEB) were cycled 98°C 1min; 35 cycles 98°C 15sec, 60°C 30sec, 72° 30sec; final extension 72°C 3min. PCR reactions were loaded on a 1% agarose gel, expected bands were cut and PCR products were extracted using GeneJet Gel Extraction Kit (Thermo Scientific) and eluted in 40μl water. Cloning was performed using the FX system as described previously (Geertsma and Dutzler, 2011). Briefly, each eluted insert was mixed with 50ng pDXinit-PAC in a molar ratio of 1:5 (vector:insert) in 10μl reactions and digested with 0.5μl SapI (NEB) for 1h at 37°C. Reactions were incubated for 20min at 65°C to heat-inactivate SapI, cooled down to room temperature and constructs were ligated by addition of 1.1μl 10x T4 ligase buffer (NEB) and 0.25μl T4 ligase (NEB) and incubation for 1h at 25°C. Ligation was stopped by heat-inactivation for 20min at 65°C followed by cooling to room temperature. 2μl ligation reactions were transformed into chemically-competent *E. coli* (NEB Stable Competent) and plated on 5% sucrose-containing LB-Ampicillin, yielding pDXinit-CD4Nb-PAC, pDXinit-CD8Nb-PAC and pDXinit-CD16Nb-PAC. For cloning of CD8 hashtag phagemids, 20ng pDXinit-CD8Nb-PAC were used as template for site-directed mutagenesis (as described earlier in this section) using primers EF156 and EF157 to generate pDXinit-CD8Nb(PH-A)-PAC, primers EF158 and EF159 for pDXinit-CD8Nb(PH-B)-PAC, primers EF164 and EF165 for pDXinit-CD8Nb(PH-C)-PAC and primers EF166 and EF167 for pDXinit-CD8Nb(PH-D)-PAC. For cloning of EGFP Nb-displaying phagemids, the EGFP Nb sequence from pOPINE GFP nanobody (Addgene ID: 49172) was amplified in 50μl PCR reactions with Q5 (NEB) using 25ng plasmid template and EF05 and EF06 primers. The EGFP nanobody insert was cloned into pDXinit using FX cloning (described earlier), yielding pDXinit-EGFPNb. EGFP Nb-displaying phagemids containing RD1 in different orientations were cloned by using pDXinit-EGFPNb and performing site-directed mutagenesis (described earlier) with EF73 and EF74 to obtain pDXinit-EGFPNb-PAC or using EF75 and EF76 yielding pDXinit-EGFPNb-RD1(5-3). For introduction of a PCR handle required for PDT library amplification, pDXinit-EGFPNb-PAC was subjected to site-directed mutagenesis (as described earlier in this section) using primers EF78 and EF79, yielding pDXinit-EGFPNb(handle)-PAC. For cloning of mCherry Nb-displaying phagemids, the mCherry Nb sequence from pGex6P1 mCherry nanobody (Addgene ID: 70696) was amplified in 50μl PCR reactions with Q5 (NEB) using 25ng plasmid template and EF07 and EF08 primers. The mCherry nanobody insert was cloned into pDXinit using FX cloning (as described earlier in this section), yielding pDXinit-mCherryNb. All constructs are listed in Supplementary Table 2.

### Analysis of RD1-mediated phagemid amplification using RD1-containing primers

5ng of either pDXinit-EGFPNb, pDXinit-EGFPNb-PAC or pDXinit-EGFPNb-RD1(5-3) were subjected to linear PCR (10μl reaction volume) using primer EF170 and 5μl 2x KAPA HiFi HotStart ReadyMix (Roche) and cycling conditions 98°C 2min; 12cycles 98°C 10sec, 59°C 30sec, 72°C 1min; final extension 72°C 5°min. After completion, 0.625μl of each primer EF147 and EF57, 1.25μl water and 12.5μl 2x KAPA were added. Nb-specific PCR was performed using 98°C 3min; 30cycles 98°C 15sec, 65°C 20sec, 72°C 1min; final extension 72°C 5min. PCR using primers EF57 and EF58 and indicated plasmid templates was used as amplification control.

### Phage production

Phagemid-containing SS320 (Lucigen) cultures were incubated overnight in 2YT medium containing 2% glucose, 50μg/ml Ampicillin and 10μg/ml Tetracycline (2YT/2%/A/T) at 37°C, 240rpm. Cultures were diluted 1:50 in 2YT/2%/A/T and grown for 2-3h at 37°C, 240rpm until OD600 = 0.4-0.5. 5ml bacteria were then infected with 200μl M13K07 helper phage (NEB) and incubated for 60min at 37°C. Bacteria were collected by centrifugation and resuspended in 50ml 2YT containing 50μg/ml Ampicillin and 25μg/ml Kanamycin (2YT/A/K). Phages were produced overnight by incubation at 37°C, 240rpm. Cultures were centrifuged and phages were precipitated from supernatants by addition of 1/4th volume 20% PEG-6000/2.5M NaCl solution and incubation on ice for 75min. Phages were collected by centrifugation (17min, 12500g, 4°C). Phage pellets were resuspended in 1.2ml PBS, suspensions were cleared (5min, 12500g, 25°C) and supernatants containing phages were stored.

### Cell culture

NIH3T3 and HEK293T cells (ATCC) were maintained in DMEM containing 10% FBS, 2 mM L-glutamine and 100 U/ml penicillin/streptomycin (Thermo Scientific) and cultured at 37°C and 5% CO_2_. For sub-culturing, medium was aspirated, cells were washed with PBS and detached with Trypsin-EDTA 0.25% (Thermo Scientific). Detachment reactions were stopped with culture medium and cells were seeded at desired densities. Cell stocks were prepared by resuspending cell aliquots in FBS with 10% DMSO and freezing them slowly at −80°C. Frozen aliquots were then moved to liquid nitrogen for long-term storage. All cell lines were regularly tested for mycoplasma contamination.

### Plasmid transfection of HEK293T cells

One day before transfection, 2×10^6^ HEK293T cells were seeded in 10cm dishes (Corning) in complete culture medium (as described in section ‘Cell culture’). Transfection was performed using GeneJuice reagent (Fisher Scientific). 600μl Opti-MEM and 12μl GeneJuice were mixed in 1.5ml tubes, vortexed shortly and spun down. 4μg of plasmid DNA (either pCAG (Addgene ID: 11160), pCAC-EGFP (Addgene ID: 89684) or pCAC-EGFP (Addgene ID: 32601)) were added, tubes were vortexed shortly and spun down. Transfection mix was added dropwise to HEK293T cells. Cells were grown for 24h at 37°C and 5% CO_2_ to allow transgene expression. Successful transfection was assessed by fluorescence microscopy on an EVOS M5000.

### Flow cytometry for detection of phage binding

Harvested cell lines or thawed PBMCs (see PHAGE-ATAC workflow for harvest and thawing protocol) were resuspended in FC buffer (above) and incubated with respective phage nanobodies for 20min on a rotator at 4°C. Cells were centrifuged and washed with cold FC buffer twice to remove unbound phages (all centrifugation steps were 350g, 4min, 4°C). For optimization of fixation and lysis conditions, cells were fixed using indicated formaldehyde concentrations (Thermo Scientific) and permeabilized with depicted lysis buffers. Cells were resuspended in FC buffer and anti-M13 antibody (Sino Biological, 11973-MM05T-50) was added at 1:500 dilution. After 10min on ice, cells were washed twice in FC buffer and anti-mouse Fc Alexa Fluor 647-conjugated secondary antibody (Thermo Scientific, A-21236) was added at 1:500 dilution. Cells were incubated for 10min on ice, washed twice in FC buffer and resuspended in Sytox Blue (Thermo Scientific) containing FC buffer for live/dead discrimination according to manufacturer’s instructions. In indicated cases, cells were stained with anti-CD4-FITC (clone OKT4, BioLegend) at 1:500 dilution, hereby no anti-M13 and anti-mouse Fc antibodies were used. Stained cells were analyzed using a CytoFLEX LX Flow Cytometer (Beckman Coulter) at the Broad Institute Flow Cytometry Facility. Flow cytometry data were analyzed using FlowJo software v.10.6.1.

### PHAGE-ATAC workflow

For cell line “species mixing” experiment, culture medium was aspirated, cell lines were washed with PBS, harvested using Trypsin-EDTA 0.25% (Thermo Scientific), resuspended in DMEM containing 10% FBS, centrifuged, washed with PBS and resuspended in FC buffer. For PBMC and CD8 T cell experiments, cryopreserved PBMCs or CD8 T cells (AllCells) were thawed, washed in PBS and resuspended in cold Flow cytometry buffer (FC buffer; PBS containing 2% FBS). All centrifugation steps were carried out at 350g, 4min, 4°C unless stated otherwise.

Cells were incubated with phages on a rotating wheel for 20min at 4°C. After three washes in FC buffer, cells were fixed in PBS containing 1% formaldehyde (Thermo Scientific) for 10min at room temperature. Fixation was quenched by addition of 2.5M glycine to a final concentration of 0.125M. Cells were washed twice in FC buffer and permeabilized using lysis buffer (10mM Tris-HCl pH 7.5, 10mM NaCl, 3mM MgCl2, 0.1% NP-40, 1% BSA) for 3min on ice. This buffer was used, as we found that standard 10x Genomics scATAC lysis buffer results in loss of pNb cell staining (**Supp. Fig. 9**). After lysis, cells were washed by addition of 1ml cold wash buffer (lysis buffer without NP-40), inverted and centrifuged (5min, 500g, 4°C). Supernatant was aspirated and the cell pellet was resuspended in 1x Nuclei Dilution Buffer (10x Genomics). Cell aliquots were mixed with Trypan Blue and counting was performed using a Countess II FL Automated Cell Counter. Processing of cells for tagmentation, loading of 10x Genomics chips and droplet encapsulation via the 10x Genomics Chromium controller microfluidics instrument was performed according to Chromium Single Cell ATAC Solution protocol.

For species-mixing, a single 10x channel was ‘super-loaded’ with 20,000 cells. Linear amplification and droplet-based indexing were performed as described in the 10x ATAC protocol on a C1000 Touch Thermal cycler with 96-Deep Well Reaction Module (BioRad). After linear PCR, droplet emulsions were broken, barcoded products were purified using MyONE silane bead cleanup and eluted in 40μl elution buffer I (Chromium Single Cell ATAC Solution protocol). At this point eluates were split for PDT and ATAC library preparation. Whereas 5μl eluate were used for PDT library preparation as described below, the remaining 35μl eluate were used for scATAC library generation (according to Chromium Single Cell ATAC Solution protocol). Splitting samples at this point is not expected to result in a loss of library complexity as PDTs and ATAC fragments already underwent amplification via linear PCR.

The aliquot for PDT library preparation was used for PDT-specific PCR in a 100μl reaction using 2x KAPA polymerase and primers EF147 and EF91, cycling conditions were: 95°C 3min, 20cycles 95°C 20sec, 60°C 30sec, 72° 20sec; final extension 72°C 5min. Amplified PDT products were purified by addition of 65μl SPRIselect beads (Beckman Coulter), 160μl supernatants were saved and incubated with 192μl SPRIselect. Beads were washed twice with 800μl 80% ethanol and the PDT library was eluted in 40μl buffer EB (Qiagen).

Concentration of PDT libraries was determined and 15ng were used for 100μl indexing PCR reactions using 50μl Amp-Mix (10x Genomics), 7.5μl SI-PCR Primer B (10x Genomics) and 2.5μl i7 sample index-containing primers (10x Genomics), cycling conditions were: 98°C 45sec; 6cycles 98°C 20sec, 67°C 30sec, 72° 20sec; final extension 72°C 1min. Indexed PDT libraries were purified by addition of 120μl SPRIselect and eluted in 40μl buffer EB. The concentration of final libraries was determined using a Qubit dsDNA HS Assay kit (Invitrogen) and size distribution was examined by running a High Sensitivity DNA chip on a Bioanalyzer 2100 system (Agilent).

PDT and ATAC libraries were pooled and paired-end sequenced (2 x 34 cycles) using Nextseq High Output Cartridge kits on a Nextseq 550 machine (Illumina). Raw sequencing data were demultiplexed with CellRanger-ATAC mkfastq. ATAC fastqs were used for alignment to the GRCh38 or mm10 reference genomes using CellRanger-ATAC count version 1.0.

### Computational workflow for generation of PDT count matrices

PDT fastqs were obtained by running CellRanger-ATAC mkfastq on raw sequencing data and custom UNIX code was used to derive PDT-cell barcode count tables. For each lane, using ‘grep -B1’ function, PDT_R3 fastqs were searched for each CDR3 barcode sequence (**Supplementary Table 4**) and corresponding sequencing cluster information was derived. Cluster information was used to derive corresponding cell barcodes from PDT_R2 fastqs by using ‘fgrep -A1 -f’. Files containing identified cell barcodes from all four lanes were concatenated, the reverse complement of cell barcode sequences was generated using ‘tr ACGTacgt TGCAtgca’ and barcodes were filtered via ‘fgrep -f’ using the cell barcodes called by CellRanger-ATAC count. Unique cell barcode occurrences were counted.

### Analysis of species mixing PHAGE-ATAC experiment

PHAGE-ATAC sequencing data from the species-mixing experiment was demultiplexed using CellRanger-ATAC mkfastq and generated ATAC fastqs were processed with CellRanger-ATAC count to filter reads, trim adapters, align reads to both GRCh38 and mm10 reference genomes, count barcodes, identify transposase cut sites, detect accessible chromatin peaks and to identify cutoffs for cell barcode calling. The “force-cells” parameter was not set. Barcodes were classified as human or mouse if >90% of barcode-associated fragments aligned to GRCh38 or mm10, respectively. Cutoffs for cell barcode calling were >3,000 ATAC fragments overlapping peaks for human and >10,000 for mouse barcodes (based on empirical density). Doublet barcodes were defined as containing more than 10% ATAC fragments aligning to both GRCh38 and mm10 reference genomes. The EGFP PDT count table was generated as described above by searching PDT fastqs for the corresponding phage barcode (**Supplementary Table 4**) and deriving PDT-associated cell barcodes via filtering using the entire list of called cell barcodes (human and mouse).

After flow cytometry measurement of HEK293T-EGFP-GPI (EGFP^+^) and HEK293T cells (EGFP^-^), FCS files were exported using CytExpert Software (Beckman Coulter). Values for forward scatter (FSC area) and EGFP fluorescence (FITC area) were derived from FCS files. Human EGFP^+^ and EGFP^-^ cells were defined based on the distribution of EGFP PDT counts (for PHAGE-ATAC) or EGFP fluorescence represented by FITC-area values (for flow cytometry) by setting a gate at the minimum value in-between both populations.

### Analysis of PBMC PHAGE-ATAC experiment

Sequencing data from two libraries of PBMCs were processed using CellRanger-ATAC count to the GRChg38 reference genome using all default parameters, yielding 7,792 high-quality PBMCs (no filtering was applied beyond the CellRanger-ATAC knee call). We downloaded processed CITE-seq PBMC data (Stoeckius et al., 2017) from the Gene Expression Omnibus (GSE100866). After removing spiked-in mouse cells, this published dataset was jointly analyzed with the 7,972 PBMCs profiled by PHAGE-ATAC. We performed data integration using canonical correlation analysis (Butler et al., 2018), using the 2,000 most variable RNA genes as is the default in Seurat. Next, we performed RNA imputation for the ATAC-seq data using Seurat v3 with the default settings (Stuart et al., 2019). Reduced dimensions and cell clusters were inferred using this merged object via the first 20 canonical correlation components with the default Louvain clustering in Seurat v3. Centered log ratio (CLR) normalized PDTs were visualized in the reduced dimension space and a per-tag, per-cluster mean was further computed to further access staining efficiency between the modalities (**Figure 1N**).

Cell annotations were derived based on well-established marker genes for PBMCs (**Supp. Fig. 10A**), and the granulocyte population was corroborated by high overall fragments but low proportion of fragments overlapping chromatin accessibility peaks. For protein-based clustering and analyses, we identified T-cell clusters from the integrated embedding (using the chromatin/RNA data) and then further stratified into subpopulations based on the abundance of the CD4 and CD8 CLR PDT abundances (**Supp. Fig. 10B**). Differential gene activity scores between these populations were then computed using the default functionality in Seurat/Signac (Wilcoxon rank-sum test).

### Analysis of cell hashing PHAGE-ATAC experiment

One channel of sequencing data from the hashed, combined CD8-enriched T cells was processed using CellRanger-ATAC count via the GRCh38 reference genome using all default parameters, yielding 8,366 high-quality PBMCs (no filtering was applied beyond the CellRanger-ATAC knee call). As we suspected the presence of contaminating B-cells, we first characterized cell states using latent semantic indexing (LSI)-based clustering and dimensionality reduction using Signac and Seurat (Stuart et al., 2019). Specifically, all detected peaks were used as input into LSI. The first 20 LSI components (except for the first component, which was found to be correlated with the per-cell sequencing depth) were used to define cell clusters using the default Louvain clustering algorithm in Seurat. Per-cluster chromatin accessibility tracks were computed using a per million fragments abundance for each cluster, as previously implemented (Lareau et al., 2020). These chromatin accessibility tracks were used to annotate cell clusters based on promoter accessibility of known marker genes.

To assign hash identities to cell barcodes, we utilized the HTODemux function from Seurat (Stoeckius et al., 2018) with the positive.quantile parameter set at 0.98. This yielded 703 doublets, 1,225 negatives, and 6,438 singlets based on the abundance and distribution of CD8 hashtag PDTs.

To verify PHAGE-ATAC hashtag-based assignments, we performed mitochondrial DNA genotyping using mgatk (Lareau et al., 2020) and nuclear genotyping and donor assignment using souporcell (Heaton et al., 2020) with “--min_alt 8 --min_ref 8 --no_umi True -k 4 --skip_remap True --ignore True” options, which resulted in 92.9% accuracy (99.3% singlet accuracy, 74% overlap in called doublets), confirming the concordance of our hashing design.

### Cloning of PANL, a synthetic high-complexity phage nanobody library

To generate randomized library inserts, three separate primer mixes (for long CDR3, medium CDR3 and short CDR3 inserts) were used for PCR-mediated assembly. For short CDR3-inserts, the primer mix contained 0.5μl each of polyacrylamide gel electrophoresis-purified EF42, EF43, EF64, EF44, EF65, EF45, EF46, EF47, EF66 and EF48 (each 100μM) (EllaBiotech). For medium CDR3-inserts, EF67 was used instead of EF66. For long CDR3-inserts, EF68 was used instead of EF66. Primer mixes were diluted 1:25 and 1μl of each mix was used for overlap-extension PCR using Phusion (NEB). Four 50μl reactions for each mix were performed using cycling conditions 98°C 1min; 20cycles 98°C 15sec, 60°C 30sec, 72° 30sec; final extension 72°C 5min. PCR reactions of the same mix were pooled and purified by addition of 280μl AMPure XP beads (Beckman Coulter). Beads were washed twice with 800μl 80% ethanol and assembled inserts were eluted in 100μl water. Concentrations of each insert (long, medium, short) were determined and pooled in a 1:2:1 molar ratio. Five identical 50μl PCR reactions with pooled inserts and primers EF40 and EF41 were performed using Phusion (NEB), cycling conditions were 98°C 1min; 30cycles 98°C 15sec, 62°C 30sec, 72° 30sec; final extension 72°C 5min. Amplified library insert was pooled and purified by adding 350μl AMPure XP beads (Beckman Coulter). Beads were washed twice with 1ml 80% ethanol and library insert was eluted in 60μl water. Five identical 60μl restriction digest reactions for digest of 7.5 μg library vector pDXinit-PAC with 2.5μl SapI were performed. Library insert (4.8μg) was digested in a 30μl reaction using 2.5μl SapI. Digests were incubated for 4h at 37°C and loaded on 1% agarose gels. Bands corresponding to digested library vector and insert were cut and products were extracted using GeneJet Gel Extraction Kit (Thermo Scientific) and eluted in 40μl water. Five identical 100μl ligation reactions were performed, each containing 1.25 μg digested pDXinit-PAC, 450ng digested insert and 0.5μl T4 ligase (NEB). Ligations were incubated for 16h at 16°C, heat-inactivated for 20min at 65°C and cooled to room temperature. 100μl AMPure XP beads were added to each ligation reaction, beads were washed twice using 300μl 80% ethanol and ligation products were eluted in 15μl water and pooled. Five electroporations in 2mm cuvettes (BioRad) were performed, each using 90μl electro-competent SS320 *E. coli* (Lucigen) and 12μl ligation product. Pulsing was performed on a GenePulserXcell instrument (BioRad) with parameters 2.5kV, 200Ohm, 25μF. After electroporation, bacterial suspensions were added to 120ml pre-warmed SOC and incubated for 30min, 37°C, 225rpm. An aliquot of library-carrying bacteria was saved at this point and used to prepare a dilution series. Each dilution was plated on LB-Ampicillin plates. After overnight incubation at 37°C, colonies were counted, transformation efficiency was determined and library complexity was estimated. The remaining 120ml of library-containing culture were added to 1.125L 2YT/2%/A/T and incubated overnight at 37°C, 240rpm. The library-containing culture was harvested, glycerol stocks were prepared and library aliquots were stored.

### Analysis of picked PANL clones using PCR and Sanger sequencing

Library-containing bacteria were plated on LB-Ampicillin, incubated overnight, and colonies were picked and inoculated in 8ml LB-Ampicillin. Cultures were incubated for at least 8h at 37°C, 240rpm. Bacteria were harvested and plasmids isolated using GeneJet Plasmid Miniprep kit (Thermo Scientific). PCR was performed to evaluate clone inserts. 10μl PCR reactions were set up that contained 10ng of isolated plasmid, 0.5μl each of primers EF52 and EF53, and 4.5μl 2x OneTaq Quick Load Master Mix (NEB). Cycling conditions were 94°C 4min; 28cycles 94°C 15sec, 62°C 15sec, 68°C 30sec; final extension 68C 5min. PCR reactions were analyzed on 2% agarose gels. Selected clones were analyzed by Sanger Sequencing using primer EF17. Observed amino acid frequencies at hypervariable positions were assessed by analyzing Sanger sequences of 25 picked clones.

### Phage nanobody library production

A PANL aliquot corresponding to 3×10^10^ bacterial cells (around 5x coverage of the library) was transferred to 200ml 2YT/2%/A/T and cultures were grown until OD600=0.5 was reached (~2h). Cultures were infected with 8ml M13K07 helper (NEB) for 60min at 37°C. Cultures were harvested, supernatants discarded and bacterial pellets were resuspended in 1L 2YT/A/K. Cultures were incubated overnight at 37°C, 250rpm for production of the input library of phage nanobody particles. Bacterial cultures were harvested, supernatants collected and phages were precipitated using PEG/NaCl as described earlier. Final phage pellets were resuspended in a total of 20ml PBS and stored. Phage titers were determined by infecting a log-phase culture of SS320 with a dilution series of the produced phage library and plating bacteria on LB-Ampicillin. Colonies were counted and titers were calculated. Produced phage libraries were characterized by titers >4×10^11^ pfu/ml.

### Phage display selection

HEK293T cells were transfected either with pCAG or pCAG-EGFP-GPI as described above. Cells were harvested, 10^7^ pCAG-transfected cells were resuspended in 1ml PBS containing 2% BSA (PBS-BSA), and 8ml PANL library (1.6×10^12^ pfu) in PBS-BSA were added for counter-selection. Samples were incubated for 1h on a rotating wheel at 4°C and then centrifuged at 350g, 5min, 4°C. Supernatants containing phages were added to 10^7^ pCAG-EGFP-GPI expressing cells for positive selection. After 1h on a rotating wheel at 4°C, samples were centrifuged (350g, 5min, 4°C) and washed 6 times with PBS-BSA to remove unbound phages. Cells were washed once in PBS, centrifuged and cell pellets were resuspended in 500μl Trypsin solution (1mg/ml Trypsin (Sigma Aldrich) in PBS) to elute bound phages. Cells were incubated for 30min on a rotating wheel at room temperature and digests were stopped by addition of AEBSF protease inhibitor (Sigma Aldrich) to a final concentration of 0.5mg/ml. Samples were centrifuged (400g, 4min at room temperature) and the supernatant containing eluted phages was used to infect 10ml of log-phase SS320 (OD600=0.4). After infection for 40min at 37°C, cultures were added to 90ml 2YT/2%/A/T and incubated overnight at 37°C, 250rpm. Cultures containing output libraries were aliquoted and glycerol stocks were prepared. Output library phage particles were prepared as described earlier for PANL and used in subsequent selection rounds using the same protocol described here.

## Acknowledgements

We thank L. Gaffney for assistance with figure illustrations and preparation, C. de Boer and other members of the Regev laboratory for helpful discussion. We acknowledge support from the Broad Institute Flow Cytometry Core facility. This research was supported by NHGRI grants 5RM1 HG006193 (Center for Cell Circuits), a gift from the Food Allergy Science Initiative, a gift from the Manton Foundation, and HHMI (to AR). E.F. is supported by an EMBO Long-Term fellowship. C.A.L. is supported by a Stanford Science Fellowship. A.R. was a Howard Hughes Medical Institute Investigator (until July 31, 2020).

## Author Contributions

E.F. conceived and designed the project with guidance from A.R. E.F. designed and performed experiments. E.F. developed the PHAGE-ATAC computational workflow with input from C.A.L. E.F. developed the PHAGE-ATAC experimental protocol with input from L.S.L. E.F. and C.A.L analyzed the data. G.E. contributed to data analysis. A.R. provided project oversight and acquired funding. E.F. and A.R. wrote the manuscript with input from all authors.

## Corresponding authors

Correspondence to E.F. and A.R.

## References

Butler, A., Hoffman, P., Smibert, P., Papalexi, E., and Satija, R. (2018). Integrating single-cell transcriptomic data across different conditions, technologies, and species. Nat Biotechnol 36, 411–420.

Gebauer, M., and Skerra, A. (2009). Engineered protein scaffolds as next-generation antibody therapeutics. Curr Opin Chem Biol 13, 245–255.

Geertsma, E.R., and Dutzler, R. (2011). A versatile and efficient high-throughput cloning tool for structural biology. Biochemistry 50, 3272–3278.

Gehring, J., Hwee Park, J., Chen, S., Thomson, M., and Pachter, L. (2020). Highly multiplexed single-cell RNA-seq by DNA oligonucleotide tagging of cellular proteins. Nat Biotechnol 38, 35–38.

Heaton, H., Talman, A.M., Knights, A., Imaz, M., Gaffney, D.J., Durbin, R., Hemberg, M., and Lawniczak, M.K.N. (2020). Souporcell: robust clustering of single-cell RNA-seq data by genotype without reference genotypes. Nat Methods 17, 615–620.

Hoogenboom, H.R. (2005). Selecting and screening recombinant antibody libraries. Nat Biotechnol 23, 1105–1116.

Ingram, J.R., Schmidt, F.I., and Ploegh, H.L. (2018). Exploiting Nanobodies’ Singular Traits. Annu Rev Immunol 36, 695–715.

Katzenelenbogen, Y., Sheban, F., Yalin, A., Yofe, I., Svetlichnyy, D., Jaitin, D.A., Bornstein, C., Moshe, A., Keren-Shaul, H., Cohen, M., et al. (2020). Coupled scRNA-Seq and Intracellular Protein Activity Reveal an Immunosuppressive Role of TREM2 in Cancer. Cell 182, 872–885 e819.

Klein, A.M., Mazutis, L., Akartuna, I., Tallapragada, N., Veres, A., Li, V., Peshkin, L., Weitz, D.A., and Kirschner, M.W. (2015). Droplet barcoding for single-cell transcriptomics applied to embryonic stem cells. Cell 161, 1187–1201.

Kubala, M.H., Kovtun, O., Alexandrov, K., and Collins, B.M. (2010). Structural and thermodynamic analysis of the GFP:GFP-nanobody complex. Protein Sci 19, 2389–2401.

Lareau, C.A., Duarte, F.M., Chew, J.G., Kartha, V.K., Burkett, Z.D., Kohlway, A.S., Pokholok, D., Aryee, M.J., Steemers, F.J., Lebofsky, R., et al. (2019). Droplet-based combinatorial indexing for massive-scale single-cell chromatin accessibility. Nat Biotechnol 37, 916–924.

Lareau, C.A., Ludwig, L.S., Muus, C., Gohil, S.H., Zhao, T., Chiang, Z., Pelka, K., Verboon, J.M., Luo, W., Christian, E., et al. (2020). Massively parallel single-cell mitochondrial DNA genotyping and chromatin profiling. Nat Biotechnol.

Ludwig, L.S., Lareau, C.A., Ulirsch, J.C., Christian, E., Muus, C., Li, L.H., Pelka, K., Ge, W., Oren, Y., Brack, A., et al. (2019). Lineage Tracing in Humans Enabled by Mitochondrial Mutations and Single-Cell Genomics. Cell 176, 1325–1339 e1322.

Ma, A., McDermaid, A., Xu, J., Chang, Y., and Ma, Q. (2020). Integrative Methods and Practical Challenges for Single-Cell Multi-omics. Trends Biotechnol 38, 1007–1022.

Macosko, E.Z., Basu, A., Satija, R., Nemesh, J., Shekhar, K., Goldman, M., Tirosh, I., Bialas, A.R., Kamitaki, N., Martersteck, E.M., et al. (2015). Highly Parallel Genome-wide Expression Profiling of Individual Cells Using Nanoliter Droplets. Cell 161, 1202–1214.

McGinnis, C.S., Patterson, D.M., Winkler, J., Conrad, D.N., Hein, M.Y., Srivastava, V., Hu, J.L., Murrow, L.M., Weissman, J.S., Werb, Z., et al. (2019). MULTI-seq: sample multiplexing for single-cell RNA sequencing using lipid-tagged indices. Nat Methods 16, 619–626.

McMahon, C., Baier, A.S., Pascolutti, R., Wegrecki, M., Zheng, S., Ong, J.X., Erlandson, S.C., Hilger, D., Rasmussen, S.G.F., Ring, A.M., et al. (2018). Yeast surface display platform for rapid discovery of conformationally selective nanobodies. Nat Struct Mol Biol 25, 289–296.

Miersch, S., and Sidhu, S.S. (2012). Synthetic antibodies: concepts, potential and practical considerations. Methods 57, 486–498.

Mimitou, E.P., Cheng, A., Montalbano, A., Hao, S., Stoeckius, M., Legut, M., Roush, T., Herrera, A., Papalexi, E., Ouyang, Z., et al. (2019). Multiplexed detection of proteins, transcriptomes, clonotypes and CRISPR perturbations in single cells. Nat Methods 16, 409–412.

Paul, F., Arkin, Y., Giladi, A., Jaitin, D.A., Kenigsberg, E., Keren-Shaul, H., Winter, D., Lara-Astiaso, D., Gury, M., Weiner, A., et al. (2015). Transcriptional Heterogeneity and Lineage Commitment in Myeloid Progenitors. Cell 163, 1663–1677.

Peterson, V.M., Zhang, K.X., Kumar, N., Wong, J., Li, L., Wilson, D.C., Moore, R., McClanahan, T.K., Sadekova, S., and Klappenbach, J.A. (2017). Multiplexed quantification of proteins and transcripts in single cells. Nat Biotechnol 35, 936–939.

Pollock, S.B., Hu, A., Mou, Y., Martinko, A.J., Julien, O., Hornsby, M., Ploder, L., Adams, J.J., Geng, H., Muschen, M., et al. (2018). Highly multiplexed and quantitative cell-surface protein profiling using genetically barcoded antibodies. Proc Natl Acad Sci U S A 115, 2836–2841.

Roobrouck, A., Stortelers, C., Vanlandschoot, P., Staelens, S., Conde, M., Soares, H., and Schols, D. (2016). Bispecific Nanobodies. US 2016/0251440 A1.

Rothbauer, U., Zolghadr, K., Tillib, S., Nowak, D., Schermelleh, L., Gahl, A., Backmann, N., Conrath, K., Muyldermans, S., Cardoso, M.C., et al. (2006). Targeting and tracing antigens in live cells with fluorescent nanobodies. Nat Methods 3, 887–889.

Satpathy, A.T., Granja, J.M., Yost, K.E., Qi, Y., Meschi, F., McDermott, G.P., Olsen, B.N., Mumbach, M.R., Pierce, S.E., Corces, M.R., et al. (2019). Massively parallel single-cell chromatin landscapes of human immune cell development and intratumoral T cell exhaustion. Nat Biotechnol 37, 925–936.

Smith, G.P. (1985). Filamentous fusion phage: novel expression vectors that display cloned antigens on the virion surface. Science 228, 1315–1317.

Stoeckius, M., Hafemeister, C., Stephenson, W., Houck-Loomis, B., Chattopadhyay, P.K., Swerdlow, H., Satija, R., and Smibert, P. (2017). Simultaneous epitope and transcriptome measurement in single cells. Nat Methods 14, 865–868.

Stoeckius, M., Zheng, S., Houck-Loomis, B., Hao, S., Yeung, B.Z., Mauck, W.M., 3rd, Smibert, P., and Satija, R. (2018). Cell Hashing with barcoded antibodies enables multiplexing and doublet detection for single cell genomics. Genome Biol 19, 224.

Stuart, T., Butler, A., Hoffman, P., Hafemeister, C., Papalexi, E., Mauck, W.M., 3rd, Hao, Y., Stoeckius, M., Smibert, P., and Satija, R. (2019). Comprehensive Integration of Single-Cell Data. Cell 177, 1888–1902 e1821.

Tavernier, J., Cauwels, A., Kley, N., and Gerlo, S. (2017). CD8 Binding Agents. WO 2017/134306 Al.

